# Regulation of the *Erythrobacter litoralis* DSM 8509 general stress response by visible light

**DOI:** 10.1101/641647

**Authors:** Aretha Fiebig, Lydia M. Varesio, Xiomarie Alejandro Navarreto, Sean Crosson

**Affiliations:** Department of Biochemistry and Molecular Biology, The University of Chicago, Chicago, IL 60637; The Committee on Microbiology, The University of Chicago, Chicago, IL 60637

**Keywords:** general stress response, HWE / HisKA2 / HisKA_2 kinase, anoxygenic aerobic photoheterotroph, Alphaproteobacteria, LOV domain

## Abstract

Extracytoplasmic function (ECF) sigma factors are a major class of environmentally-responsive transcriptional regulators. In *Alphaproteobacteria* the ECF sigma factor, σ^EcfG^, activates general stress response (GSR) transcription and protects cells from multiple stressors. A phosphorylation-dependent protein partner switching mechanism, involving HWE/HisKA_2-family histidine kinases, underlies σ^EcfG^ activation. The identity of these sensor kinases and the signals that regulate them remain largely uncharacterized. We have developed the aerobic anoxygenic photoheterotrophic (AAP) bacterium, *Erythrobacter litoralis* DSM 8509, as a comparative genetic model to investigate GSR regulation. Using this system, we sought to define the contribution of visible light and a photosensory HWE kinase, LovK, to GSR transcription. We identified three HWE kinases that collectively regulate GSR: *gsrK* and *lovK* are activators, while *gsrP* is a repressor. GSR transcription is higher in the dark than light, and the opposing activities of *gsrK* and *gsrP* are sufficient to achieve light-dependent differential transcription. In the absence of *gsrK* and *gsrP*, *lovK* alone is sufficient to regulate GSR transcription in response to light. This regulation requires a photochemically active LOV domain in LovK. Our studies establish a role for visible light and HWE kinases in light-dependent regulation of GSR transcription in *E. litoralis*, an AAP species.

**GRAPHICAL ABSTRACT:** 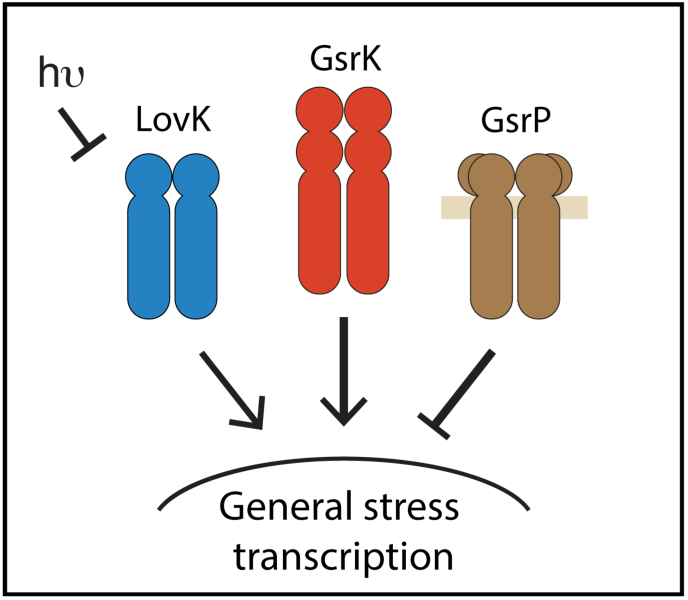

**ABBREVIATED SUMMARY:** General stress response (GSR) systems protect bacteria from a diverse range of physical and chemical stressors. We have developed *Erythrobacter litoralis* as a new genetic model to study GSR in *Alphaproteobacteria* and show that three HWE-family histidine kinases collectively regulate GSR transcription via σ^EcfG^. Visible light is a GSR regulatory signal in *E. litoralis*, and LovK is a blue-light photosensor kinase that functions as a dark activated GSR regulator.

## INTRODUCTION

### LOV-HWE kinases: An overview

Light is a ubiquitous environmental factor that provides energy to support life in many ecosystems. Multiple adaptive responses to light have evolved in bacteria including phototaxis and photoavoidance, photoprotective pigment production, and the regulation of genes required for photosynthesis. These responses are mediated by proteins that sense photons in particular energy ranges across the visible spectrum (Purcell & Crosson, 2008). LOV domains are widely distributed photosensors (Losi *et al*., 2002) that detect blue light via a bound flavin cofactor (Christie *et al*., 1999). These photosensory domains are present in an assortment of signal transduction proteins from bacteria, archaea, fungi, protists, and plants (Glantz *et al*., 2016, Herrou & Crosson, 2011). Though broadly conserved (Losi *et al*., 2015, Villar *et al*., 2018), the physiological roles of LOV photosensors in bacteria remain largely undefined.

The class *Alphaproteobacteria* contains species that inhabit diverse ecological niches, and that have a significant impact on nutrient cycling, agricultural production and human health (Batut *et al*., 2004). *Alphaproteobacteria* commonly encode proteins that contain a LOV domain coupled to an HWE/HisKA2-type (Herrou *et al*., 2017) histidine kinase. Surprisingly, these “LOV-HWE kinases” are often present in heterotrophic species with no evident photobiology (Herrou & Crosson, 2011). Histidine kinases are typically co-expressed with and phosphorylate their cognate response regulators to directly control gene expression (Hoch & Silhavy, 1995), but HWE/HisKA2 kinases are unusual in that they are often orphaned on the bacterial chromosome, or are adjacent to single domain response regulators (SDRR) that lack a regulatory output domain to control gene expression (Herrou *et al*., 2017).

Early studies of alphaproteobacterial LOV-HWE kinases in *Caulobacter crescentus* (Purcell *et al*., 2007) and *Rhizobium leguminosarum* (Bonomi *et al*., 2012), demonstrated their influence on cell adhesion and biofilm formation. In the case of *C. crescentus*, a LOV-HWE kinase (LovK) modulates adhesion by controlling expression of a single downstream gene that regulates surface adhesin biosynthesis (Fiebig *et al*., 2014, Reyes Ruiz *et al*., 2019). However, the major effect of deleting or overexpressing genes encoding LOV-HWE kinases appears to be dysregulation of the general stress response (GSR) system (Foreman *et al*., 2012, Kim *et al*., 2014), which determines cell survival across a range of stress conditions (Fiebig *et al*., 2015, Francez-Charlot *et al*., 2015). There is increasing evidence that multiple HWE/HisKA2-family kinases function as part of complex regulatory networks that regulate the GSR in *Alphaproteobacteria* by influencing the phosphorylation state of the anti-anti-σ factor, PhyR (Kaczmarczyk *et al*., 2014, Gottschlich *et al*., 2018, Lori *et al*., 2018, Correa *et al*., 2013). Phospho-PhyR activates an extracytoplasmic function (ECF) σ factor – EcfG – by binding and sequestering its anti-σ factor, NepR (Francez-Charlot *et al*., 2009) (Figure 1).

**Figure 1:**
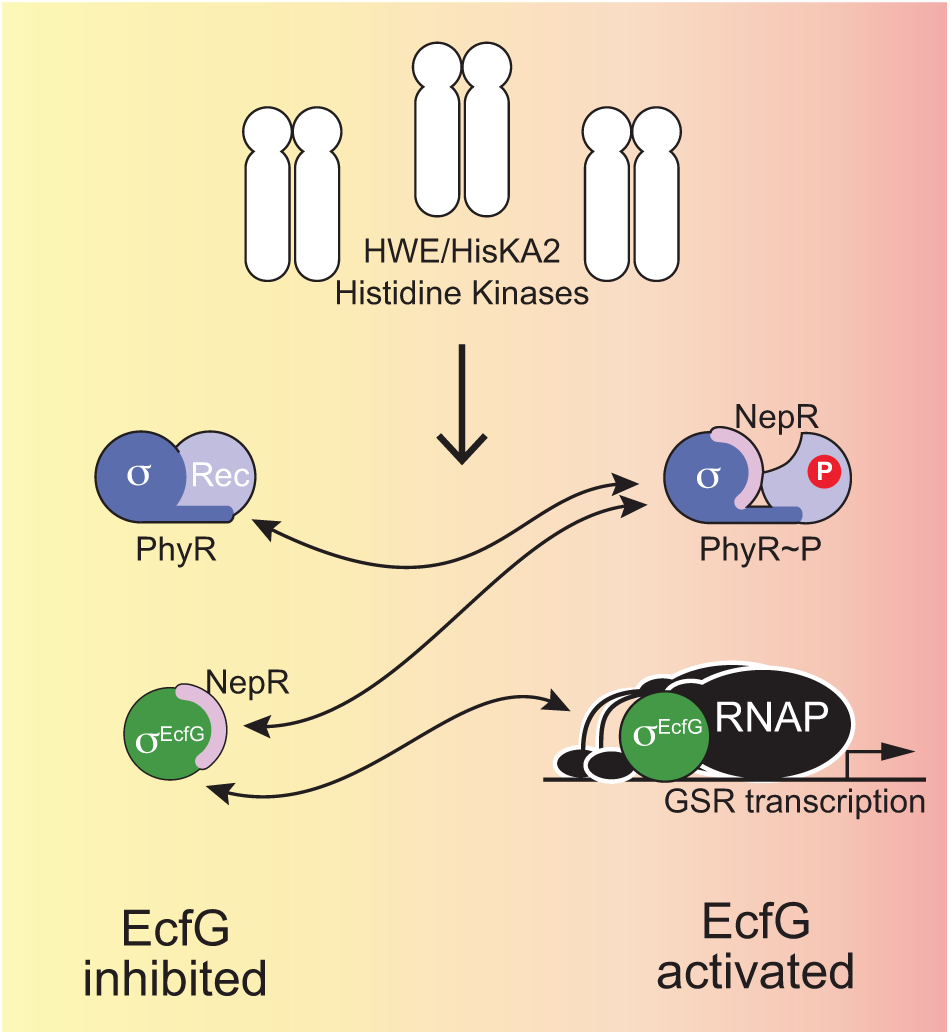
Core regulators of the alphaproteobacterial general stress response. In *Alphaproteobacteria*, the general stress response (GSR) is often controlled by an ensemble of HWE/HisKA_2 family sensor histidine kinases that modulate the phosphorylation state of PhyR. PhyR phosphorylation promotes a protein partner switch that sequesters the anti-sigma factor, NepR, thereby releasing the extracytoplasmic function (ECF) sigma factor, EcfG, to initiate transcription of genes in the general stress response regulon. The core protein partner switch genes, *phyR*, *nepR* and *ecfG* are broadly conserved in *Alphaproteobacteria*. The number of kinases controlling the pathway is variable.

Though there is genetic evidence that LOV-HWE kinases control transcription of genes in the GSR regulon (Foreman *et al*., 2012, Kaczmarczyk *et al*., 2014, Kim *et al*., 2014, Sycz *et al*., 2015), the effects of visible light on LOV-HWE kinase signaling *in vivo* remain undefined. Therefore, it is important to identify appropriate experimental systems to assess the effect of visible light and other environmental signals on LOV-HWE kinase signaling in living cells.

### Erythrobacter litoralis DSM 8509 as a system to study LOV-HWE kinase signaling

We sought to investigate the signaling role of LOV-HWE kinases in the genus *Erythrobacter*, which was attractive to us for several reasons. *Erythrobacter* spp. are abundant in the world’s oceans (Sunagawa *et al*., 2015, Tully *et al*., 2018, Villar *et al*., 2018) where they contribute to global nutrient and energy cycling (Koblizek *et al*., 2007, Kolber *et al*., 2001). Importantly, LOV-HWE kinases are common in *Erythrobacter* spp., as evidenced in both sequenced isolates (Oh *et al*., 2009, Wang *et al*., 2014) and in a set of metagenome assembled genomes (MAGs) (Delmont *et al*., 2018) from the Tara oceans sequence datasets (Villar *et al*., 2018). Though *Erythrobacter* spp. can be isolated and grown in axenic culture (Lei *et al*., 2015, Wang *et al*., 2014), strains have not been extensively cultivated and manipulated. Thus, *Erythrobacter* spp. are less likely to have acquired lab-adaptive mutations that could potentially skew the effects of light on cell physiology. Most *Erythrobacter* spp. are aerobic anoxygenic photoheterotrophs (AAP), which carry a photosynthesis gene cluster that encodes production of bacteriochlorophyll *a* (Bchl *a*), and other genes required for phototrophy. Therefore, physiological responses to visible light are expected in this genus.

*Erythrobacter litoralis* strain DSM 8509 was isolated from a cyanobacterial mat in the sublittoral zone off the island of Texel, Netherlands (Yurkov *et al*., 1994, Yurkov & Van Gemerden, 1993). Draft genome sequence of DSM 8509 indicated the presence of a single LOV-HWE kinase gene (Wang *et al*., 2014). This distinguishes DSM 8509 from *E. litoralis* strain HTCC 2594, which encodes multiple LOV-kinases (Oh *et al*., 2009) that have been previously characterized *in vitro* (Correa *et al*., 2013, Dikiy *et al*., 2019, Rivera-Cancel *et al*., 2014). HTCC 2594 does not encode the genes required for phototrophy (Oh *et al*., 2009, Zheng *et al*., 2016), and though DSM 8509 and HTCC 2594 cluster phylogenetically based on their 16S sequences, they do not group based on amino acid sequence of concatenated core genes (Zheng *et al*., 2016). As such, it has been suggested that HTCC 2594 should be re-classified as a distinct species, *Erythrobacter* sp. HTCC 2594 (Jiang *et al*., 2018).

Here, we report the development of *E. litoralis* DSM 8509 as an experimental genetic system, which we have used to study the contribution of a LOV-HWE kinase to regulation of the general stress response. Our data show that an ensemble of three HWE kinases, including the LOV-HWE kinase LovK, collectively regulates the general stress response in *E. litoralis*. The cytoplasmic kinase, GsrK, functions as an activator of GSR transcription in complex medium, while the transmembrane kinase, GsrP, is a repressor. LovK activates GSR transcription, and is a more potent GSR activator under dark conditions. Transcription of the entire GSR regulon is significantly higher in dark conditions than in light. In fact, the set of genes that are differentially transcribed upon shifts in the light environment strongly overlaps the experimentally-defined GSR regulon. While LovK contributes to light-dependent regulation of GSR transcription, it is not strictly required for this response. Our results support a model in which photons directly and indirectly modulate GSR transcription via the HWE kinases GsrK, GsrP, and LovK.

## RESULTS

### A brief comparison of *Erythrobacter litoralis* DSM 8509 to other *Erythrobacter* species

Several partial and complete genome sequences of *Erythrobacter* isolates have been deposited in public databases (see Table 1 for representative genomes). These genomes have similar characteristics in terms of size and GC content, yet differ in presence/absence of particular genes and pathways. For example, *Erythrobacter* spp. can encode zero, one, or multiple LOV-HWE kinases, and the total number of HWE/HisKA2-family kinases ranges from two to sixteen (Ulrich & Zhulin, 2010). Although the three LOV kinases present in strain HTCC 2594 have been biochemically characterized (Correa *et al*., 2013, Dikiy *et al*., 2019, Rivera-Cancel *et al*., 2014), we anticipated that the potential functional redundancy of these genes could complicate interpretation of genetic and physiological studies *in vivo*. In contrast, DSM 8509 encodes only one LOV-HWE kinase. Additionally, when considering our intent to investigate the interplay between multiple HWE/HisKA2-family kinases involved in GSR regulation, we noted that DSM 8509 has a ‘goldilocks’ number of such kinases: more than one, but not too many to complicate genetic analyses (Table 1). Finally, DSM 8509 encodes core genes required for phototrophy, which provides an opportunity to develop genetic tools in a species for which light is likely a central environmental/metabolic signal. For these reasons, we chose to pursue the development of *E. litoralis* DSM 8509 as a comparative model to study the function of LOV-HWE kinases.

**Table 1:** Genome characteristics of select *Erythrobacter* spp. isolates.

| Isolate <sup>a</sup> | Source (reference) | Genome status | Genome length (Mb) | % GC | Phototrophy genes | LOV domain <sup>b</sup> | LOV kinases (SDRR - orphan) <sup>c</sup> | HWE / HisKA2 kinases <sup>d</sup> |
| --- | --- | --- | --- | --- | --- | --- | --- | --- |
| *E. longus* DSM 6997 (Och 101) | Seaweed in Aburatsubo Bay, Kanagawa, Japan (Shiba & Simidu, 1982) | WGS; 14 contigs (Wang *et al.*, 2014) | 3.6 | 57 | YES | 0 | 0 | 2 |
| *E. litoralis* DSM 8509 | Cyanobacterial mat, Supralitoral zone, Texal Netherlands (Yurkov *et al.*, 1994, Yurkov & Van Gemerden, 1993) | WGS; 22 contigs (Wang *et al.*, 2014) | 3.21 | 65 | YES | 1 | 1-0 | 5 |
|  |  | Complete (this reference) |  |  |  |  |  |  |
| *Erythrobacter* sp. SD-21 | Surface sediment, San Diego Bay, USA (Francis *et al.*, 2001) | WGS; 19 contigs | 2.97 | 63 | NO | 1 | 0-1 | 16 |
| *Erythrobacter* sp. NAP1 | Surface water, Northwest Atlantic Ocean (Koblizek *et al.*, 2003) | WGS; 4 contigs (Koblizek *et al.*, 2011) | 3.26 | 61 | YES | 1 | 0-1 | 6 |
| *E. litoralis* HTCC2594 | 10 m deep Sargasso Sea, Atlantic Ocean (Oh *et al.*, 2009) | Complete (Oh *et al.*, 2009) | 3.05 | 63 | NO | 4 | 1-2 | 8 |

<sup>a</sup> Isolates sequenced at the time this work was initiated.
<sup>b</sup> Extracted from (Glantz *et al.*, 2016)
<sup>c</sup> Subset of genes identified in (Glantz *et al.*, 2016) with kinase domains in an operon with a single domain response regulator (SDRR) or encoded as an orphan gene. All of these are HWE/HisKA\_2-family kinases.
<sup>d</sup> Identified in the MiST database 2.0 (Ulrich & Zhulin, 2010)

### Whole-genome sequencing and development of genetic tools in DSM 8509

To our knowledge, genetic analysis of an *Erythrobacter* species has not been reported. To support development of *Erythrobacter litoralis* DSM 8509 as an experimental genetic model, we first completed and closed the genome sequence. Specifically, we re-sequenced DSM 8509 using a PacBio long-read sequencing platform to an average depth of 143X (Table S1). These sequence reads were assembled *de novo* to a single 3.25 Mb contig, which was deposited and is available to the public under NCBI GenBank accession CP017057. We surveyed the antibiotic sensitivity profile of DSM 8509 to identify useful markers for genetic selection (Table S2), and adapted methods to transform the bacterium with both replicating (pBBR-derived) and integrating (ColE1-derived) plasmids. We successfully transformed *E. litoralis* DSM 8509 by both electroporation and conjugation (see Materials and Methods). We further identified conditions to generate unmarked gene deletion and allele replacement strains using a two-step recombination and *sacB*/sucrose counterselection approach, which we applied to our genetic analysis of HWE kinases and GSR signaling described below.

### *E. litoralis* DSM 8509 LovK is a photosensor

Like *Caulobacter crescentus* (Marks *et al*., 2010, Purcell *et al*., 2007), *E. litoralis* DSM 8509 possesses a single LOV-HWE histidine kinase gene that is located adjacent to a single-domain response regulator gene on the chromosome (locus tags Ga0102493_111685 and 111686). We have named these genes *lovK* and *lovR*, respectively (Figure 2). The N-terminal LOV domain of LovK contains a complete flavin binding consensus sequence (Crosson & Moffat, 2001, Herrou & Crosson, 2011), and a conserved cysteine residue required for light-dependent cysteinyl-flavin adduct formation (Salomon *et al*., 2000). To validate that LovK indeed functions as a *bona fide* photoreceptor *in vitro*, we cloned and expressed LovK in a heterologous *E. coli* system, and purified the protein by affinity chromatography. LovK has a classic LOV domain visible absorption spectrum, with a λ_max_ at 450 nm and vibronic bands at 425 nm and 475 nm. Illumination of the protein with blue light results in loss of these major bands, and a concomitant increase in absorption at 396 nm (Figure 3). This spectral signature is consistent with formation of a covalent cysteinyl-flavin C4(a) adduct upon illumination (Crosson & Moffat, 2002, Salomon *et al*., 2000, Salomon *et al*., 2001, Swartz *et al*., 2001). We further expressed and purified LovK(C73A), in which the conserved cysteine postulated to form a flavin C4(a) adduct is mutated to a non-reactive alanine. LovK(C73A) retains flavin binding as evidenced by its absorption spectrum, but does not undergo the light-dependent bleaching of visible absorption bands that are indicative of cysteinyl-flavin adduct formation (Figure 3). These data provide evidence that *E. litoralis* DSM 8509 LovK is a photosensor that has the photochemical features of typical LOV proteins.

**Figure 2:**
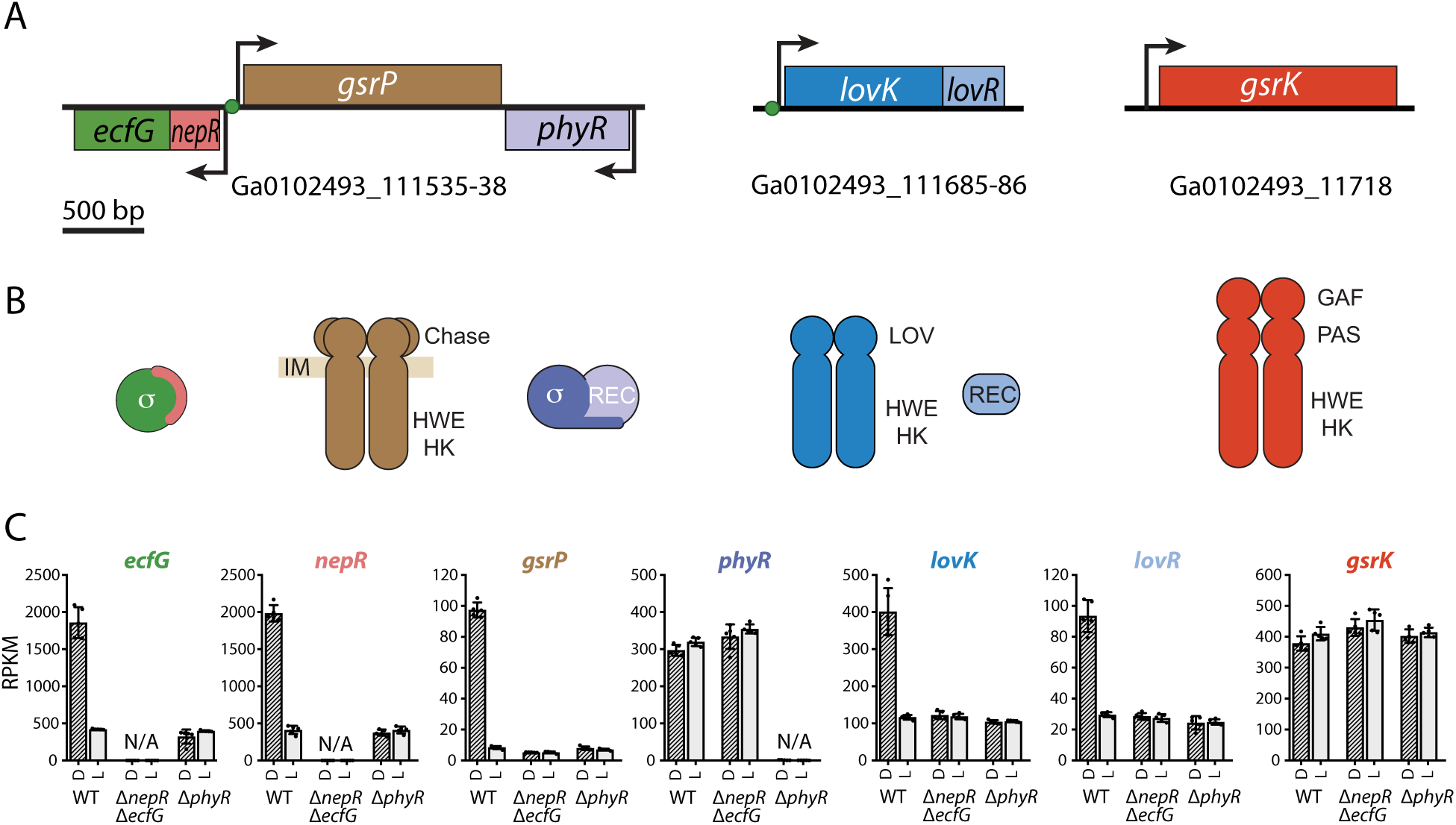
GSR regulators and their expression in *E. litoralis* DSM 8509. **A.** Genes encoding the GSR regulators in *E. litoralis*. The core partner switch genes (*phyR*, *nepR*, and *ecfG*) together with a HWE kinase gene (*gsrP*) constitute the GSR locus (left). The *lovKR* operon and *gsrK* are not adjacent to each other or to the GSR locus on the chromosome. Gene locus numbers (GenBank accession CP017057) are listed below each gene diagram. Predicted EcfG-binding motifs are marked with green circles at the base of the transcription start arrows. **B.** Domain structure of the proteins encoded by the genes in (A). Colors match the gene colors. GsrP is predicted to encode a transmembrane histidine kinase and is depicted in the inner membrane (IM). All other proteins lack transmembrane domains and are predicted to be cytoplasmic. **C.** RNA-seq analyses of transcript abundance from the genes in (A). Reads per kilobase per million reads (RPKM) of each gene is plotted for wild-type (WT), Δ*nepR-ecfG* or Δ*phyR* strains grown in the dark (D - hashed bars) or the light (L – light grey bars). Mean ± s.d. is plotted. n=5 per treatment; individual values from experimental replicates are dots.

**Figure 3:**
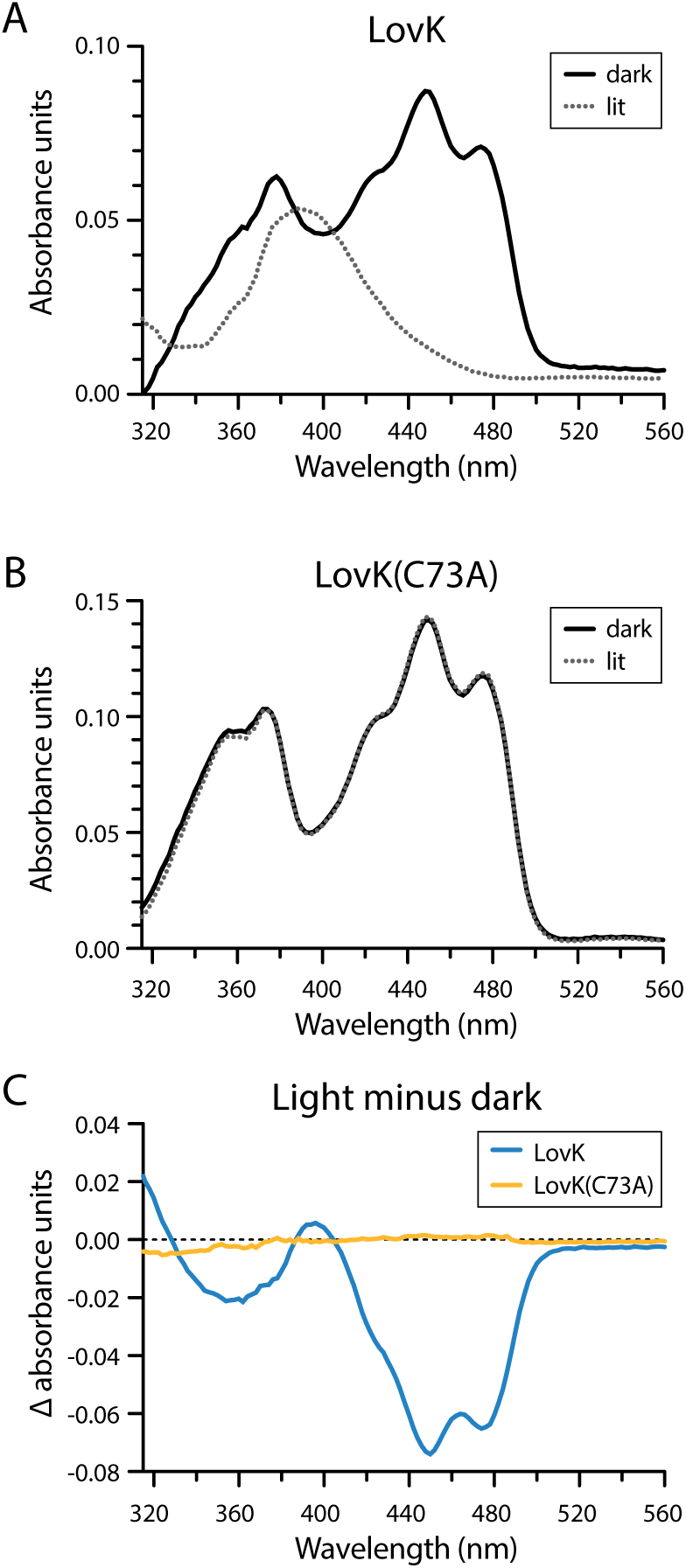
Absorption spectroscopy provides evidence that LovK protein can function as a photosensor. **A-B.** Visible absorption spectrum of purified *E. litoralis* DSM 8509 (A) LovK and (B) LovK(C73A) proteins. Lit spectrum was collected immediately after illuminating the protein with blue light. The dark spectrum is the same protein that was not illuminated. **C.** Light minus dark difference absorption spectrum for LovK (blue line) and LovK(C73A) (yellow line). Photoexcitation of the wild-type protein results in a signature loss of absorbance between at 440 and 480 nm and an increase in absorbance at 396 (Salomon *et al*., 2000, Swartz *et al*., 2001) due to cysteinyl-flavin C4(a) adduct formation (Crosson & Moffat, 2002, Salomon *et al*., 2001). The LovK(C73A) mutant protein lacks the conserved cysteine required for adduct formation and does not exhibit light induced changes in absorbance.

### The *E. litoralis* DSM 8509 GSR regulon

Before testing whether LovK plays a role in the regulation of general stress response (GSR) transcription, we first sought to define the GSR regulon in *E. litoralis* DSM 8509 by RNA-sequencing (RNA-seq). To this end, we deleted the predicted orthologs of *phyR* (locus tag Ga0102493_111538) and the *nepR-ecfG* operon (locus tags Ga0102493_111536 and 111535) (Figure 2). As discussed in the introduction, these genes encode a protein partner switching system that controls activation of GSR transcription by σ^EcfG^. Deletion of either *phyR* or *ecfG* were predicted to result in a strain incapable of activating GSR transcription. We compared the global transcriptional profiles of the Δ*phyR* and Δ*nepR-ecfG* deletion strains to wild type (WT) (Table S3). To examine the effects of light, each strain was grown in cool white fluorescent light (60 µmol m^−2^ s^−1^) or in darkness (foil-covered tubes) for 24 hours prior to harvesting mRNA.

A shared set of 183 genes were differentially expressed (more than 1.5-fold; false discovery rate (FDR) p-value < 0.01) in Δ*phyR* and Δ*nepR-ecfG* compared to wild-type cells (Figure 4, Table S4). With few exceptions, transcript levels in this gene set were lower in the two mutant strains compared to wild type indicating that these differentially regulated genes are transcriptionally activated by σ^EcfG^ in wild-type cells. We identified a predicted σ^EcfG^ binding motif that matched EcfG motifs from other *Alphaproteobacteria* (Fiebig *et al*., 2015, Francez-Charlot *et al*., 2015, Staron & Mascher, 2010, Staron *et al*., 2009) in the promoter region of approximately one-quarter of the regulated genes (Figure 4, Table S4). Presence of these motifs predict the subset of genes in the *E. litoralis* GSR regulon that are directly regulated by the ECF sigma factor, σ^EcfG^.

**Figure 4:**
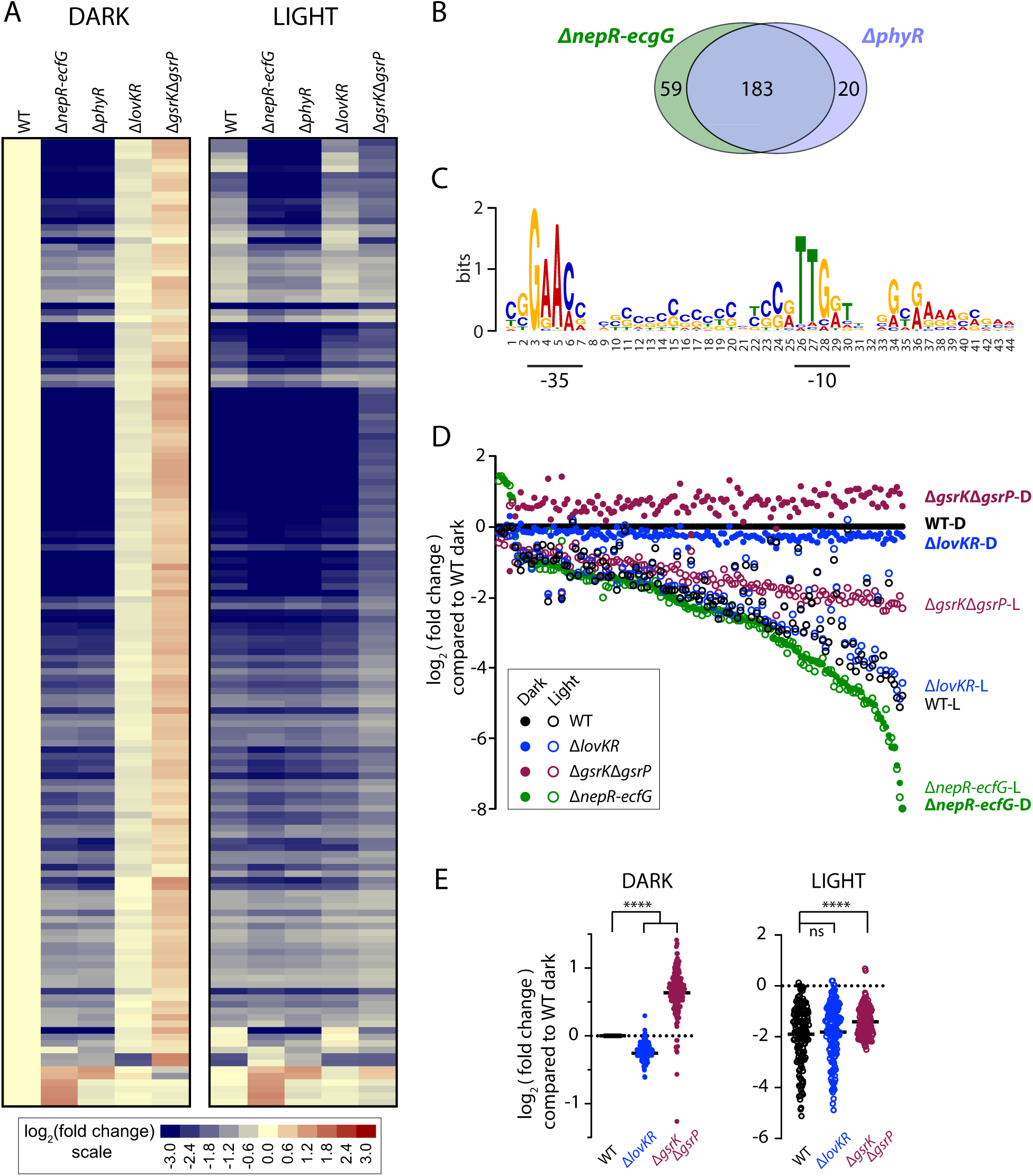
General stress response regulon of *E. litoralis* DSM 8509. **A.** Heatmap showing relative expression of genes in the GSR regulon, defined by RNA-sequencing. Each row represents a gene, and each column represents a genotype-condition pair. Wild-type (WT) and mutant strains grown in the dark are on the left and strains grown under constant white light illumination (∼60 µmol m^−2^ s^−1^) are on the right. The color or each block represents the log_2_(fold change) for an individual gene, where fold change is the ratio of RPKM_(indicated condition)_ / RPKM_(WT dark)_. Color scale is below the heat map. The heat map contains a subset of 148 genes in Table S4 with a fold change > 2 in the Δ*nepR-ecfG* or the Δ*phyR* data sets, and a max group mean RPKM > 10. GSR regulatory genes that were deleted in these strains were excluded from the gene set presented in the heatmap. **B.** Venn diagram of the differentially regulated genes in the Δ*nepR-ecfG* or Δ*phyR* strains compared to wild type. The criteria for genes deemed to be differentially regulated was a fold change > 1.5 with a false-discovery rate (FDR) p-value < 0.01. These genes in are listed in Table S4. **C.** Logo of the ECF σ-type motif identified in the promoter region of a subset of genes in the GSR regulon; the logo was generated using MEME motif discovery tools (Bailey *et al*., 2009). The −35 and −10 boxes (underlined) of this *E. litoralis* motif are consistent with previously described EcfG motifs (Staron *et al*., 2009, Francez-Charlot *et al*., 2015, Fiebig *et al*., 2015). **D.** Relative expression of the 148 genes in panel A. Each position on the x-axis represents a gene ranked (left to right) by log_2_(fold change) in Δ*nepR-ecfG* dark relative to WT dark. Color-coded dots at each position represent the relative expression of a gene in different strain-condition combinations compared to WT dark (see key). Relative expression in dark-grown cultures are closed circles; light-grown cultures are open circles. **E.** Relative expression values for genes in the GSR regulon in WT, Δ*lovKR* and Δ*gsrK*Δ*gsrP* strains (grown in light or in dark) presented in a single column. Expression of each gene in these strain backgrounds is plotted relative to WT dark. The mean difference between each column was statistically assessed by one-way ANOVA followed by Dunnett’s multiple comparison test; **** indicates p<0.0001. The key in D corresponds to both graphs in E.

Genes encoding transporters, cell envelope, and cell surface proteins are broadly represented in the *E. litoralis* DSM 8509 GSR regulon (Table S4). Transcription of genes encoding the ferritin-like protein DPS, a SOUL heme-binding protein, and superoxide dismutase (*sod*) are commonly under the control of the GSR system in other *Alphaproteobacteria* (Alvarez-Martinez *et al*., 2007, Britos *et al*., 2011, Foreman *et al*., 2012, Francez-Charlot *et al*., 2009, Gourion *et al*., 2008, Gourion *et al*., 2009, Jans *et al*., 2013, Kim *et al*., 2013, Kim *et al*., 2014, Lourenco *et al*., 2011, Martinez-Salazar *et al*., 2009, Sauviac *et al*., 2007), and are regulated by *nepR-ecfG* and *phyR* in *E. litoralis* DSM8509 (Table S4). The promoters of *dps, sodA, sodC*, and a predicted SOUL heme-binding protein gene (Ga0102493_112016) contain EcfG motifs and thus appear to be directly activated by σ^EcfG^. Our transcriptomic data also provide evidence that expression of several DNA photolyases is activated by the GSR system.

Genes directly or indirectly regulated by σ^EcfG^ that may be relevant to the photobiology of *E. litoralis* include a gene encoding a TspO/MBR tryptophan-rich protein (locus Ga0102493_112559). This class of outer membrane protein has been reported to bind tetrapyrroles and promote their photooxidative degradation (Ginter *et al*., 2013), and to negatively regulate photosynthesis in *Rhodobacter* (Yeliseev & Kaplan, 1995). Transcripts levels from this particular TspO/MBR gene are highly reduced in strains lacking *phyR* or *nepR-ecfG*. Unlike the fungus *Aspergillus fumigatus*, where TspO/MBR expression is strongly induced by light exposure (Fuller *et al*., 2013), expression of *E. litoralis* TspO/MBR (Ga0102493_112559) is lower in light than in dark conditions. Notably, steady-state transcript levels from the bacteriochlorophyll biosynthesis gene, *bchC*, are higher in both Δ*phyR* and in Δ*nepR-ecfG* relative to wild type (Table S4). Transcripts of other bacteriochlorophyll biosynthesis genes including *bchF*, *bchB* and *bchN* are also higher in Δ*phyR* and Δ*nepR-ecfG* strains, though to a lesser extent than *bchC* (Figure S1). These results indicate that the *E. litoralis* GSR system directly or indirectly represses select genes involved in phototrophy.

### Light-dependent regulation of transcription at the genome scale

In wild-type *E. litoralis*, transcripts in the GSR regulon are more abundant in dark-grown cells (aluminum foil-covered tubes) than in cells illuminated with white light (60 µmol m^−2^ s^−1^) (Figure 4). In fact, measured GSR transcript levels from illuminated wild-type cells are only slightly higher than mutant strains lacking either *phyR* or *nepR-ecfG*. From this result, we conclude that the GSR is only marginally active when cells are grown in continuous white light (Figure 4 and Table S4). A light versus dark difference in GSR transcripts is not observed in Δ*phyR* or Δ*nepR-ecfG* cells.

In wild-type cells, the majority of genes differentially expressed between light and dark conditions are in the GSR regulon. Of the 220 transcripts that change more than 1.4-fold (FDR p-value < 0.01), only 37 are not in the GSR regulon as defined (Figure 5, Table S5). Twenty-five of the differentially-expressed genes outside of the GSR regulon have similar light-dependent changes in the Δ*phyR* and Δ*nepR-ecfG* strains, and thus represent a GSR-independent light-regulated gene set. This GSR-independent gene set exhibits modest (less than 2-fold) regulation in response to white light treatment, and includes photosynthesis genes with reduced transcript levels in the light (Figure 5, Figure S1). These genes are distinct from those derepressed in the GSR mutant strains (e.g. *bchC*) and include *bchH*, *bchL*, *puhA*, and *bchM*. We conclude that there are genes involved in photosynthesis (e.g. *bch*, *puf*, *puh*) for which expression is influenced by the GSR system, and others for which regulation is independent of GSR (Figure S1).

**Figure 5:**
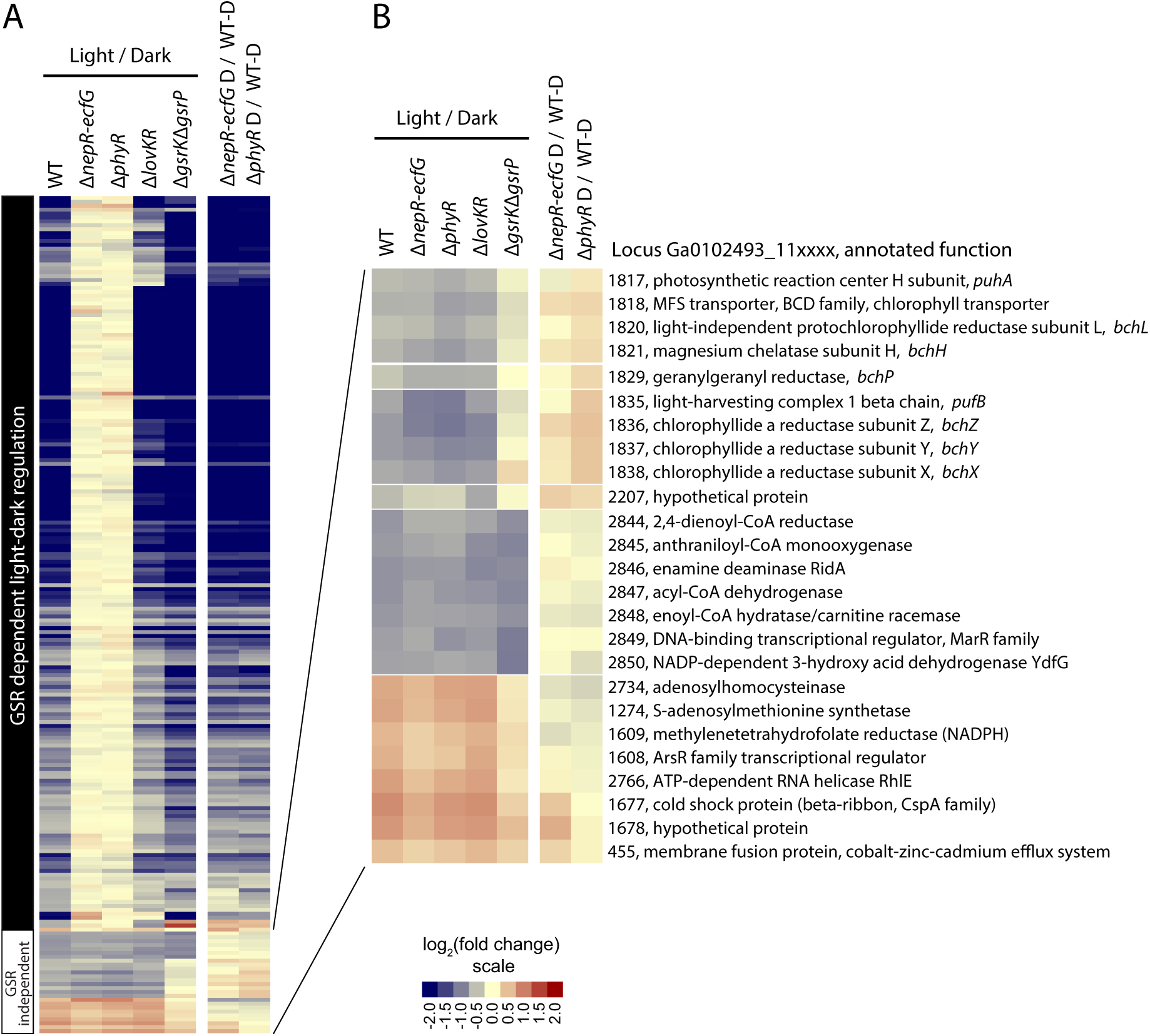
The *E. litoralis* light-dark regulon overlaps with the GSR regulon. **A.** Heatmap of differentially expressed genes in wild-type (WT) *E. litoralis* DSM 8509 grown in constant ∼60 µmol m^−2^ s^−1^ white light or in the dark (D). Cutoff criteria for differential expression are fold change > 1.4 and FDR p-value<0.01 (see Table S5 for list of genes). Heatmap includes only genes with max mean RPKM >10. For each gene (in rows), the log_2_(RPKM_light_ / RPKM_dark_) is shown for each genotype (columns). In addition, relative change in the Δ *nepR-ecfG* and Δ *phyR* strains compared to wild type (all grown in the dark) is presented to highlight congruence between light-regulated genes and the GSR regulon. **B.** Heatmap highlighting genes at the bottom of the cluster in panel A. These genes exhibit light-dependent regulation, but are not in the GSR regulon. Color scale corresponds to both panels.

Reduction in photosynthesis gene expression in *E. litoralis* in the light is consistent with light repression of reaction center genes in the purple photosynthetic bacterium, *Rhodobacter capsulatus* (Zhu & Hearst, 1986), and downregulation of photosynthesis-related genes in the aerobic anoxygenic phototrophic bacterium, *Dinoroseobacter shibae*, upon a shift from heterotrophic growth in the dark to photoheterotrophic growth in the light (Tomasch *et al*., 2011). An operon of unknown function (Ga0102493_112844-50) involved in acyl-CoA metabolism is repressed by light, independent of the GSR system. Eight metabolic genes of varying function are activated by light, independent of the GSR system (Figure 5).

### Regulators of the GSR signaling pathway

Typically, transcription of genes encoding the core GSR regulators NepR, EcfG, PhyR and GSR sensory kinases is activated by σ^ecfG^ (Foreman *et al*., 2012, Francez-Charlot *et al*., 2009, Gourion *et al*., 2009, Jans *et al*., 2013, Kim *et al*., 2013, Kim *et al*., 2014, Martinez-Salazar *et al*., 2009, Sauviac *et al*., 2007). In *E. litoralis*, a σ^EcfG^-binding motif is positioned directly upstream of the *nepR-ecfG* operon. As expected, *nepR-ecfG* transcripts were reduced in the strain lacking *phyR* and in cells grown in the light (Figure 2C, Table S4).

It was previously noted that conserved nucleotides at −35 and −10 within the σ^EcfG^-binding motif are nearly palindromic, which lead to the hypothesis that a palindromic site might function to drive bi-directional expression of oppositely oriented genes (Staron *et al*., 2009). The −35 and −10 sites upstream of *nepR-ecfG* have a strong palindromic character (GGAAC-N_17_-GTTCC) (Figure 6, Table S4). Our transcriptomic data provide evidence that expression of both *nepR-ecfG* and the oppositely oriented HWE sensor kinase gene (Ga0102493_111537, *gsrP*) is activated from this single palindromic site. Both *nepR*-*ecfG* and *gsrP* are part of the GSR regulon as determined by RNA-seq (Figure 2C, Table S4). Moreover, RNA-seq reads corresponding to *nepR-ecfG* and *gsrP* transcripts each begin 13-14 bp downstream of either end of this shared promoter motif (Figure 6). While this is not a definitive mapping of transcriptional start sites, these data strongly suggest that this particular motif can functional bi-directionally. The mechanism by which σ^EcfG^ could regulate transcription from both strands at this site is not known. We note that the motif oriented toward *nepR-ecfG* has a stronger score in MEME, and that more RNA-seq reads mapped to *nepR-ecfG* than *gsrP*, which together suggest that σ^EcfG^ preferentially initiates transcription of *nepR-ecfG*. Finally, we observed a small number of RNA-seq reads that mapped to the negative strand of this motif and its downstream region (genome positions 619,592-619,621, Figure 6). We speculate that these reads may correspond to transcription initiation from a cryptic or non-ECF σ promoter that would ensure some minimal level of *nepR-ecfG* expression under conditions in which the GSR was completely inactive.

**Figure 6:**
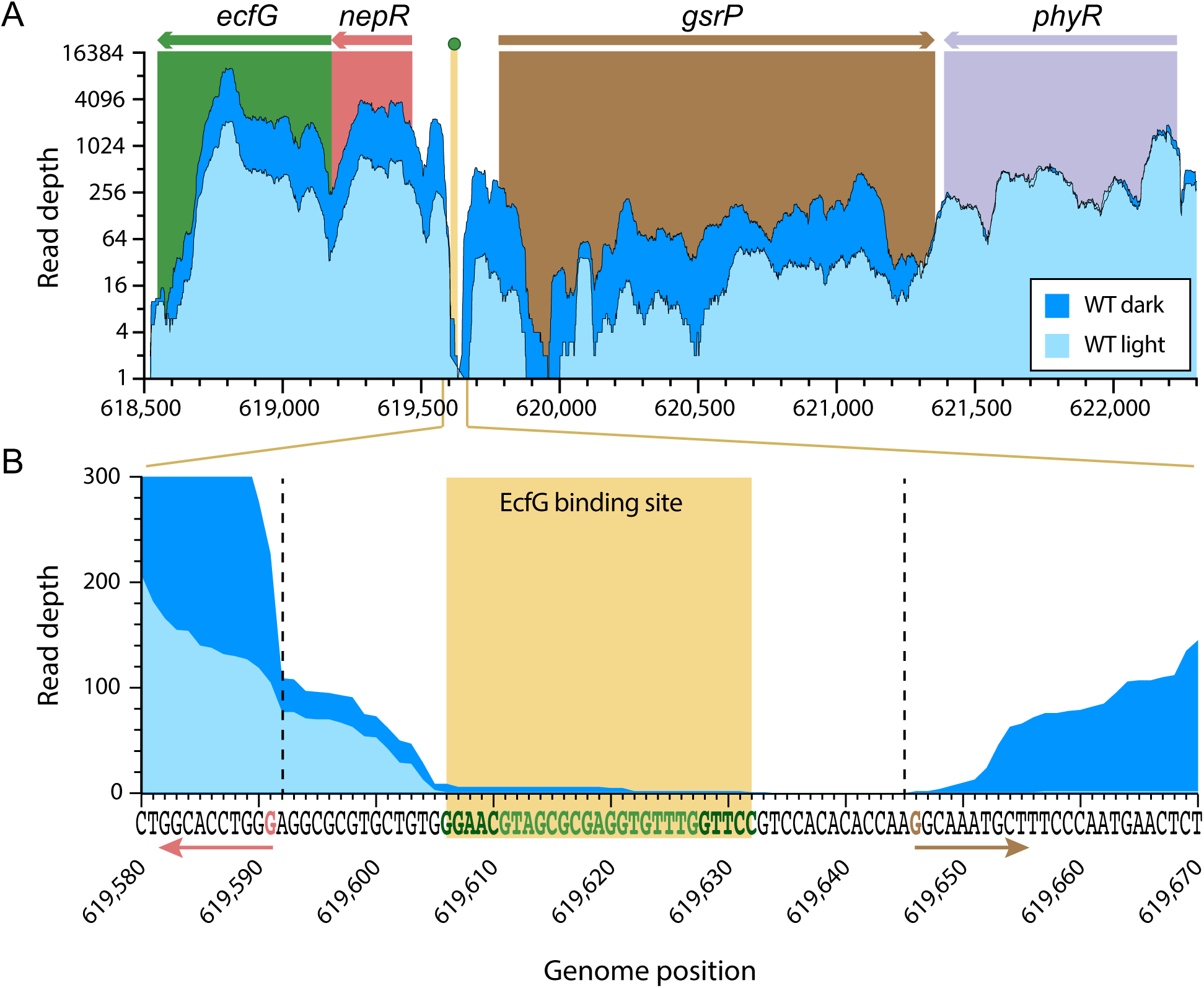
Palindromic σ^EcfG^ binding site lies between *nepR-ecfG* and *gsrP*. **A.** RNA-seq reads mapped to the GSR locus from wild-type cells grown in the dark or in the light. Number of mapped reads are plotted as a function of genome position (GenBank accession CP017057). The genes in this region are colored as in Figure 2. The single σ^EcfG^ motif in this region is marked by a green dot and beige bar. Mapped read depth for *phyR* is similar in light and dark conditions, while RNA-seq reads are more abundant in the dark for *nepR-ecfG* and *gsrP*. **B.** Expansion of the intragenic region between *nepR-ecfG* and *gsrP*. Palindromic bases of the −35 and −10 sites of EcfG motif are dark green, and the intervening bases are lighter green. The beginning of the peak of reads for each transcript is marked with a vertical dashed line. The predicted +1 sites are colored and marked with an arrow to indicate the direction of transcription (colors correspond to the genes as in panel A).

In contrast to many previously described systems, an EcfG motif was not evident in the *phyR* promoter and, accordingly, transcript levels for this gene were not affected by deletion of *nepR*-*ecfG* or by light conditions (Figure 2C and 6, Table S4). In other words, *phyR* appears to be constitutively expressed in *E. litoralis* DSM 8509. Constitutive expression of *phyR* likely necessitates the presence of a negative PhyR regulator to avoid constitutive activation of the GSR.

HWE/HisKA2-family kinases are associated with the GSR regulatory system in *Alphaproteobacteria* (Staron & Mascher, 2010, Fiebig *et al*., 2015, Francez-Charlot *et al*., 2015). *E. litoralis* DSM 8509 encodes five HWE/HisKA2-family kinases. Of these, only three are expressed at an appreciable level under our laboratory cultivation conditions, including: *1)* the kinase encoded adjacent to *nepR-ecfG* operon (Ga0102493_111537), *2) lovK*, and *3)* an orphan sensor kinase (Ga0102493_11718) (Figure 2). Ga0102493_111537 encodes a transmembrane HWE sensor kinase with a periplasmic CHASE (Mougel & Zhulin, 2001) sensory domain; its transcription is activated by *phyR* and *ecfG* and by dark conditions (Figure 2C & 6). As noted above, a shared EcfG motif identified between *nepR-ecfG* and this kinase likely promotes expression of both transcripts. The *lovK-lovR* promoter also contains an EcfG motif, and transcription of *lovK-lovR* requires *nepR-ecfG* and *phyR* (Figure 2C). The orphan HWE kinase, Ga0102493_11718, lacks an EcfG motif in its promoter and is constitutively expressed in our experimental conditions (Figure 2C). Of the remaining HWE-kinase genes, transcripts corresponding to the orphan kinase gene Ga0102493_112963 are nearly undetectable (RPKM < 10). HWE-kinase gene, Ga0102493_111751, is encoded in an operon with a CRP-family transcription factor and a single domain response regulator. Transcripts for these genes are also low abundance (Figure S2).

### The role of three HWE-family sensor kinases in regulation of GSR transcription

To assess the role of HWE-family kinases 111537, 11718, and LovK in regulation of GSR transcription, we constructed strains bearing in-frame unmarked deletions of each kinase gene. We then evaluated GSR transcriptional output in these strains by measuring levels of a GSR-regulated transcript, *dps*, by qRT-PCR. This transcript was selected because *1)* it exhibits a large dynamic range of expression between “GSR-ON” and “GSR-OFF” conditions (Figure S3), *2)* it contains an EcfG motif in its promoter suggesting direct regulation by σ^EcfG^ (Table S4), and *3)* primers to this transcript provided efficient amplification in one-step qRT-PCR reactions (see Materials and Methods). Gene locus Ga0102493_112759, which encodes a methylmalonyl-CoA mutase, was selected as a control gene for normalization (Figure S3). In our RNA-seq data sets, transcripts for this gene were abundant (in the top 20% of all transcripts) and the RPKM coefficient of variation between samples was among the lowest, making this a suitable control gene for normalization. We tested these qRT-PCR primer sets in the same strains evaluated by RNA-seq and patterns of *dps* expression were consistent between these two methods (Figure S3).

In a strain lacking kinase 111537, steady-state levels of *dps* transcript increased in cultures grown both in the light and in the dark (Figure 7A). These data are consistent with 111537 functioning as a negative regulator of GSR transcription. Conversely, deletion of the 11718 resulted in decreased *dps* transcript levels, consistent with this gene functioning as a positive regulator of GSR transcription. Both of these transcriptional phenotypes were complemented by expression of the deleted gene from its native promoter on a replicating plasmid (Figure S4). We have named these genes *gsrP* and *gsrK*, for *g*eneral *s*tress *r*esponse *p*hosphatase and *k*inase respectively. We note that the putative phosphatase and/or kinase activity of these proteins has not been established, but our genetic data are consistent with these biochemical activities against PhyR.

**Figure 7.**
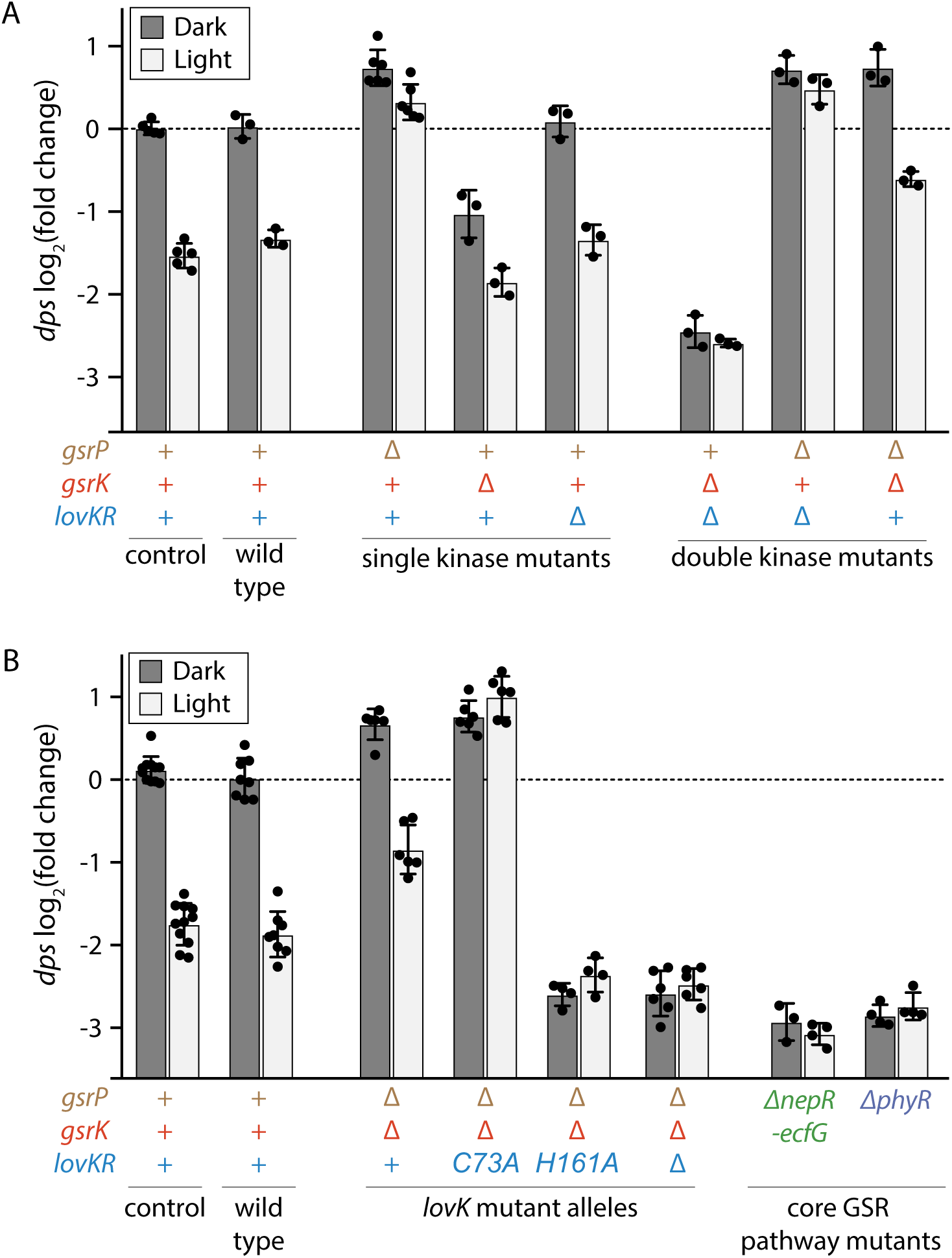
Combinatorial control of GSR transcription by three HWE-family sensor histidine kinases. **A-B.** qRT-PCR quantification of the levels of, *dps*, a GSR-regulated transcript. *dps* transcript measurements were carried out on RNA isolated from (**A**) a set of single and double HWE kinase mutant strains and (**B**) *lovK* mutant, Δ*nepR-ecfG* and Δ*phyR* strains grown in the dark (D, dark grey bars) or the light (L, light grey bars). Each measurement represents the (Ct*_dps_* – Ct_control_)_sample_ – average (Ct*_dps_* – Ct_control_)_WT-D_, which yields a log_2_(fold change) in *dps* level relative to WT dark. Strains in each panel were grown and assayed together. Each sample was assayed in triplicate. Points represent the average value for each sample. Bars represent mean ± s.d. of all samples in each condition. As an internal control, a pair of wild type-dark and wild type-light samples was assayed on every plate (control).

Deletion of the *lovKR* operon did not have a significant effect on *dps* transcript levels as measured by qRT-PCR (Figure 7A). Moreover, we did not observe large differences in GSR transcription in a Δ*lovKR* strain at the genome scale compared to wild type in our RNA-seq experiments (Figure 4). Nonetheless, GSR transcripts were uniformly lower in Δ*lovKR* compared to wild type when strains were grown in the dark (Figure 4D&E). The average log_2_(fold change) difference in all GSR transcripts between these strains is −0.25, which reflects ∼15% reduction in GSR transcription in the Δ*lovKR* strain (p < 0.0001). This trend is specific to the GSR regulon: the average log_2_(fold change) difference for all transcripts is 0.01 reflecting that, on average, the transcriptome at the whole-genome scale remains unchanged. These results provide evidence that *lovKR* plays a subtle role as an activator of GSR transcription in the dark.

To examine possible redundancy in the functions of these HWE kinases, we evaluated the effect of deleting pairs of kinases, leaving one of the three genes intact. In a Δ*gsrK* Δ*lovKR* double deletion strain where *gsrP* remains, *dps* transcript levels are lower than either of the single deletion strains in both the light and dark (Figure 7A). This result supports a model in which both GsrK and LovK function as GSR activators; moreover GsrP does not activate GSR transcription in the dark or in the light. In the Δ*gsrP* Δ*lovKR* double deletion strain where *gsrK* remains, *dps* transcripts are elevated in both the light and the dark, similar to the strain lacking only *gsrP*. Finally, in the Δ*gsrP* Δ*gsrK* strain where *lovKR* remains, *dps* transcripts are higher than wild type in both the light and the dark (Figure 7A) and are differentially expressed in response to light treatment. In this strain, *dps* levels are higher in dark-grown than light-grown cells. To further investigate whether LovK functions as a global activator of GSR, we measured the transcriptome of the Δ*gsrP* Δ*gsrK* strain grown in the light and in the dark by RNA-seq. These experiments confirm the single gene (*dps*) results in this strain. Specifically, GSR transcripts in this strain are broadly elevated compared to wild type in either light condition, and are higher in the dark than the light (Figure 4, Table S4). Together these data support a model in which LovK can function as an activator of GSR transcription that has enhanced activity in the dark.

To test whether LOV domain photochemistry (i.e. cysteinyl-flavin covalent adduct formation) contributes to the light-dark difference in LovK-dependent transcription, we mutated the conserved LOV domain cysteine (C73) to an alanine in the Δ*gsrP* Δ*gsrK* strain. Purified LovK(C73A) mutant protein is blind to light (Figure 3), and in the Δ*gsrP* Δ*gsrK lovK(C73A)* strain, GSR transcription was no longer sensitive to light (Figure 7B). Specifically, *dps* levels were comparably high in light and dark grown cultures and similar to dark grown Δ*gsrP* Δ*gsrK lovKR^+^*cultures. From these data, we again conclude that LovK is a more potent activator of GSR in its dark state.

We further tested whether the conserved histidine phosphorylation site in LovK (H161) was required to regulate GSR transcription. *dps* transcript levels in the Δ*gsrP* Δ*gsrK lovK(H161A)* strain are similar to a strain lacking all three kinases (Δ*gsrP* Δ*gsrK ΔlovKR)* indicating that a *lovK(H161A)* mutant behaves like a *ΔlovKR* deletion. We conclude that LovK phosphorylation is necessary for LovK to activate GSR transcription (Figure 7B). Together, the data provide evidence that *gsrP*, *gsrK* and *lovK* coordinately regulate GSR transcription under our assayed conditions.

## DISCUSSION

Regulation of transcription by alternative sigma factors including σ^S^ in *Gammaproteobacteria* (Battesti *et al*., 2011, Hengge, 2008), σ^B^ in select Gram-positive bacteria (Hecker *et al*., 2007), and σ^EcfG^ in *Alphaproteobacteria* (Fiebig *et al*., 2015, Francez-Charlot *et al*., 2015) confers general resistance to a range of physical and chemical stress conditions *in vitro*. Though these general stress response (GSR) σ factors are conserved within phylogenetic groups, the input signals that cue their activation and the σ-dependent transcriptional outputs vary across species. GSR inputs and outputs presumably reflect the distinct physicochemical challenges that particular species encounter within their niches. In this study, we report the development of the aerobic anoxygenic photoheterotroph (AAP), *Erythrobacter litoralis* DSM 8509, as a comparative genetic model system to study regulation of GSR transcription by σ^EcfG^. More specifically, we have sought to define the regulatory role of HWE-family sensor histidine kinases (Herrou *et al*., 2017) — including a photoresponsive LOV-HWE kinase — in σ^EcfG^-dependent transcription in *E. litoralis*.

### Light, LOV, and bacterial stress responses

Proteins containing photosensory LOV domains are widely distributed in archaea, eukarya, and bacteria (Glantz *et al*., 2016). The possibility that visible light regulates bacterial stress responses via LOV domains was first noted (Losi *et al*., 2002) after the discovery of the *Bacillus subtilis* LOV-STAS protein, YtvA, which functions as an activator of σ^B^-dependent transcription (Akbar *et al*., 2001). It was later shown that blue light can indeed activate σ^B^-dependent transcription in *B. subtilis* via YtvA (Avila-Perez *et al*., 2006). Subsequent studies of a related LOV-STAS protein from *Listeria monocytogenes* support the conclusion that light regulation of σ^B^ by LOV proteins occurs more broadly in Gram-positive bacteria (O’Donoghue *et al*., 2016, Ondrusch & Kreft, 2011). A regulatory link between LOV and σ^S^ has also been reported in *Pseudomonas syringae*, where white light was shown to repress expression of the gene encoding σ^S^ (*rpoS*) via a LOV histidine kinase (Moriconi *et al*., 2013).

Though the Alphaproteobacterial GSR sigma factor, σ^EcfG^, is not related to σ^B^ or σ^S^, studies of *Caulobacter crescentus* LovK provide evidence that LOV-HWE kinases repress σ^EcfG^-dependent transcription (Foreman *et al*., 2012). Experiments in *Brucella abortus* have identified a related LOV-HWE kinase, LovhK, that activates σ^EcfG^-dependent transcription (Kim *et al*., 2014, Sycz *et al*., 2015). However, these studies have not provided evidence that visible light is an input signal that influences σ^EcfG^-dependent transcription. In *E. litoralis* DSM 8509, we have shown that the activity LovK as a regulator of σ^EcfG^ is influenced by light. More specifically, LovK functions with two additional HWE-family kinases, GsrP and GsrK, to control GSR output (Figure 8A): GsrK is a strong GSR activator, GsrP is a repressor, and LovK is an activator that can be directly modulated by light (Figure 8A). These data contribute to an emerging model of GSR regulation in *Alphaproteobacteria* in which consortia of HWE/HisKA2-family kinases regulate σ^EcfG^.

**Figure 8:**
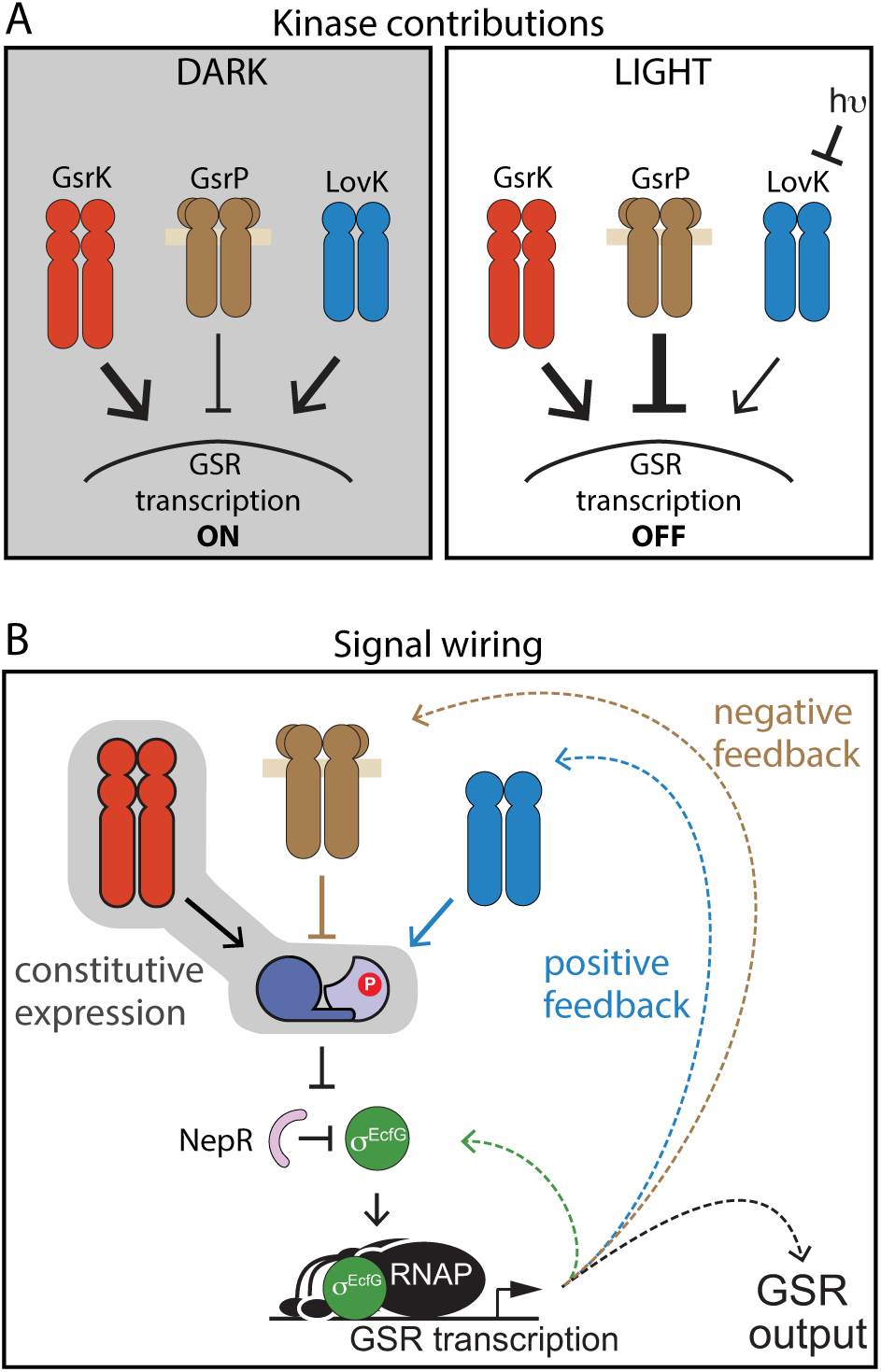
Model of GSR regulation in *E. litoralis* DSM 8509. **A.** Three HWE-family sensor histidine kinases (LovK, GsrK, and GsrP) control output of σ^EcfG^-dependent transcription. The weight of arrows shows the relative regulatory contribution of each sensor kinase under light and dark conditions. **B.** Model of GSR control network including protein interactions (solid lines) and transcriptional output (dashed lines). Most, but not all, of the GSR pathway regulators are subject to feedback regulation. PhyR, which serves to integrate signals, and GsrK, the primary activating kinase, are not under transcriptional control of σ^EcfG^, and are instead constitutively expressed. This feature of the GSR system in *E. litoralis* DSM 8509 likely necessitates a negative regulator to prevent constitutive activation of GSR transcription; GsrP fulfills this function.

Our RNA-seq experiments show that *E. litoralis* GSR transcription is highly activated in the dark, and only marginally activated in cells continuously illuminated with white fluorescent light. In contrast, light transiently induces transcription of the core GSR regulators (Dshi_3834-3837) and other stress response genes in the related AAP species, *Dinoroseobacter shibae* (Tomasch *et al*., 2011). Similarly, blue light activates *rpoE*- and *rpoH*-dependent stress responses in the phototrophic alphaproteobacterium *Rhodobacter sphaeroides* (Braatsch *et al*., 2004, Ziegelhoffer & Donohue, 2009). We did not measure transient responses of *E. litoralis* to light or dark shifts, and the advantage (if any) of elevated GSR transcription in the dark is not known.

### LovK functions as part of a consortium of GSR sensor histidine kinases

Although LovK can clearly function as a photoreceptor, it is not explicitly required for light/dark control of GSR transcription. Specifically, transcription from a GSR-regulated promoter remains light-responsive in a *gsrK*^+^ *gsrP*^+^ Δ*lovKR* strain. Thus, the combined activities of GsrK and GsrP must be balanced in a manner that leads to higher transcriptional output in dark relative to illuminated conditions (Figure 7A). This light responsiveness could emerge from enhanced activity of GsrK in the dark, enhanced repression by GsrP in the light, or a combination of both. There is no evidence to suggest that photons are a direct signal for either sensor, thus we predict that either GsrK and/or GsrP indirectly sense a metabolic or physiological change that occurs upon shifts in the light environment.

We favor a model in which GsrP is a more potent GSR repressor in cells grown in the light (Figure 8A). In our experimental conditions, *gsrK* alone strongly activates GSR transcription in light and dark (Figure 7A). This gene is constitutively expressed in both conditions (Figure 2C). While our results suggest that GsrK activity is not affected by light, we cannot entirely exclude the possibility that light conditions influence GsrK. GSR output from the strain bearing only *gsrP* is equivalently low in light and dark conditions (Figure 7A). While light may influence the activity of GsrP as a repressor, it is difficult to assess this possibility in the absence of a kinase that activates GSR transcription. σ^EcfG^–dependent transcription of *gsrP* comprises a negative feedback loop that controls GSR transcriptional output; the level of *gsrP* transcripts is dramatically reduced in light conditions (Figure 2C, Table S4). This regulatory circuitry leads one to predict that the activity of GsrP as a repressor is inversely proportional to its steady-state levels in the cell. The activities of *E. litoralis* LovK, GsrK, and GsrP likely serve to both counteract and reinforce each other depending on environmental conditions. The possibility that *E. litoralis* and other *Alphaproteobacteria* can coordinately perceive multiple environmental inputs through a consortium of HWE-family kinases may endow these species with the ability to execute complex decision-making processes with regard to control of GSR transcription.

Finally, we note that our data support a model in which LovK is active in both the light and dark, though activity of LovK as a positive regulator of σ^EcfG^-dependent transcription is enhanced in the dark. These *in vivo* data are consistent with *in vitro* data showing that light does not regulate *E. litoralis* HTCC 2594 LOV kinases in a binary (i.e. ON-OFF) manner, but rather modulates phosphorylation kinetics (Correa *et al*., 2013). Though blue light typically enhances the enzymatic activity of LOV kinases *in vitro* (Correa *et al*., 2013, Swartz *et al*., 2007), there is precedent for the unlit, dark state being the active state *in vivo*. In *D. shibae*, a short LOV protein is a dark activator of pigment biosynthesis (Endres *et al*., 2015). In this case, the LOV domain may be thought of as a dark sensor rather than a light sensor.

One may speculate on reasons why LOV proteins are often incorporated into GSR systems in the bacterial kingdom, but the fact remains that we have little understanding of the physiological relevance of LOV-regulated responses to changes in the light or redox environment in bacteria. *E. litoralis* DSM 8509 is a tractable experimental system with interesting physiological features. This species can now be leveraged as a comparative model to investigate core molecular mechanisms underlying environmental stress responses in *Alphaproteobacteria*.

## MATERIALS AND METHODS

### Growth of *E. litoralis*

*Erythrobacter litoralis* DSM 8509 was obtained from the American Type Culture Collection (ATCC 700002). For these studies, this organism was grown in Difco Marine Broth 2216. When broth was prepared according to manufacturer’s instructions, cells tended to clump in flocs. Dilution of marine broth to 0.5X reduced flocculation, and thus all cultures were grown at this broth dilution (18.7 g of powder per L). 0.5X marine broth was sterilized by autoclaving and, prior to inoculation with *E. litoralis*, the medium was passed through a sterile 0.22 µm filter to remove precipitates. Liquid cultures were grown in an Infors air incubator at 30°C in glass culture tubes, inclined at a 45° angle, shaking at 200 RPM. For “light” conditions, a bay of 10 fluorescent Philips and Sylvania T8 lights inside the shaking incubator was switched on. For “dark” conditions glass tubes were carefully wrapped in aluminum foil. For molecular genetic manipulation and isolation of mutants, colonies were grown on 0.5X marine broth solidified with 15 g of agar per L of medium. Agar plates were grown in a 30°C air incubator, or at room temperature in ambient light.

### Growth of *Escherichia coli* strains

*E. coli* strains were grown in LB Miller medium (10 g peptone, 5 g yeast extract, 10 g NaCl per L) at 37°C. Growth medium was solidified with 1.5% agar. Antibiotics were added at the following concentrations as appropriate: kanamycin, 50 µg/ml; gentamycin 15 µg/ml; chloramphenicol 12 µg/ml.

### Antibiotic sensitivity testing

To evaluate the appropriate antibiotic concentrations to use for selection in genetic manipulations, *E. litoralis* DSM 8509 cells were first grown in a streak on 0.5X marine agar at 30°C for 2-3 days. Cells were scraped from the agar plate into 1 ml of 0.5X marine broth and evenly suspended by pipetting up and down. The optical density (OD) at 660 nm was was adjusted to ∼ 0.1 AU. Cells were then 10-fold serially diluted and 20 µl of each dilution was spotted onto 0.5X marine broth agar that had been supplemented with a range of antibiotics concentrations. The antibiotic concentrations tested were as follows: 200, 100, 50, 25, 10 µg/ml for ampicillin, carbenicillin and tetracycline; 100, 50, 20, 10, 5, 1 µg/ml for gentamycin, spectinomycin, streptomycin, apramycin, and rifampicin. After the liquid absorbed into the agar, the plates were incubated at 30°C for 1 week. The minimal antibiotic concentrations that resulted in at least 4 orders of growth inhibition are reported in Table S2.

### Molecular cloning for plasmid generation

Plasmids for expression or allele replacement were generated using routine techniques. Genes or loci of interest were PCR amplified with KOD Xtreme Hot Start polymerase (Millipore Sigma) using the primers listed in Table S6. To aid amplification of these GC-rich sequences, PCR reactions were supplemented with 5% DMSO (final concentration). Primers included a 5’ extension with restriction endonuclease sites or overhangs for overlap extension PCR reactions. Null alleles were generated by first amplifying fragments ∼500 bp upstream and ∼500 bp downstream of the gene of interest. These two fragments were “stitched” together with overlap extension PCR to generate the desired allele. PCR products were cleaned using GeneJet DNA cleanup columns (Thermo Fisher), digested with appropriate restriction endonucleases (NEB) to generate overhang sequences, and ligated into similarly digested plasmids using T4 DNA ligase (NEB). Ligation reactions were transformed into chemically competent TOP10 *E. coli* by heat shock. Transformants were selected on LB supplemented with the appropriate antibiotic and grown overnight at 37°C. The inserted sequence and cloning junctions of all plasmids were confirmed by PCR amplification with plasmid specific primer sequences, treatment with ExoSapIT (Applied Biosystems) followed by Sanger sequencing (University of Chicago Comprehensive Cancer DNA Sequencing and Genotyping Facility). Plasmids generated for and used in this study are listed in Table S6.

### Transformation of *E. litoralis*

We evaluated and optimized two methods for introducing DNA into *E. litoralis* DSM8509, electroporation and conjugation. Replicating plasmids (with BBR or RK2 replication origins) could be introduced by electroporation, but this method was not efficient enough to introduce non-replicating (i.e. suicide) plasmids for chromosomal integration. Conjugation, by triparental mating, was used to introduce suicide plasmids containing *oriT* sequences, and also replicating plasmids containing *oriT* sequences. Gentamycin (10 µg ml^−1^) was used to select for clones carrying pBVMCS-4 derived plasmids. Chloramphenicol (1 µg ml^−1^) was used to select for clones carrying integrated plasmids, which permitted us to generate in-frame deletion or allele replacement strains. Detailed electroporation and conjugation protocols follow.

#### Electroporation

Cells were scraped from a freshly grown plate that had been grown for 3 days at 30°C. A pellet of 150-200 µl of cells was suspended in 750 µl 0.5x marine broth. Cells were pelleted by centrifugation (1 min at 14,000 × g) at 4°C. The cell pellet was washed 3 times with 750 µl ice cold sterile water with a 1 min centrifugation step at each wash. After the final wash, the cells were resuspended with 140 µl cold water. The total volume of cells and water was ∼200 µl. 60 µl cells were mixed with 0.1-1 µg purified plasmid and placed in a 1 mm electroporation cuvette. Cells were subjected to a single 1.8 kV pulse using the EC1 setting on a MicroPulser (Bio-Rad). Time constants were ∼ 4-5 msec. 450 µl of 0.5x marine broth was added to the cuvette and cells were transferred in this broth to a sterile 13 mm glass culture tube and incubated at 30°C shaking at 200 rpm for 2-4 hours. After this outgrowth, cells were plated on selective medium (about 200 µl per 100 mm petri dish). These plates were incubated at 30°C for about 1 week. After 5 days, pinprick-sized colonies were visible. After 7-8 days, colonies were picked and struck on fresh selective medium to confirm antibiotic resistance and to select away any non-transformed cells. We confirmed that the clones carried the plasmid of interest by colony PCR, amplifying plasmid specific sequences using cells as the template.

#### Conjugation

Plasmids encoding *oriT* sequences were transferred from *E. coli* TOP10 donor strains to E*. litoralis* using a helper strain (MT607 / pRK600) (Finan *et al*., 1986) which encodes the pilus required to mobilize *oriT* containing sequences. *E. litoralis* recipient strains were inoculated from fresh plates into 2 ml 0.5x marine broth in 13 × 100 mm glass culture tubes and grown shaking at 30°C overnight. The *E. coli* TOP10 donor strain carrying the plasmid to be transferred was grown in 2 ml LB supplemented with the appropriate antibiotic and the *E. coli* helper strain was grown in 2 ml LB supplemented with chloramphenicol at 37°C overnight. The three cultures were mixed at a ratio of 1 ml recipient, 200 µl donor, 200 µl helper. The mixed cells were centrifuged for 1 min at 10,000-14,000 × g. The pellet was resuspended in 50-100 µl 0.5x marine broth and then spotted onto a fresh 0.5x marine broth agar plate. After the liquid in the spot of cells absorbed into the agar, the plate was incubated overnight at room temperature or 30°C. The mix of cells was scraped from this non-selective plate and spread on 0.5x marine broth agar supplemented with *1)* the appropriate antibiotic to select for acquisition of the plasmid and *2)* 100 µg /ml nalidixic acid (Nal) to counterselect against the *E. coli* donor and helper strains. *E. coli* strains exhibit increased resistance to Nal when grown on marine broth agar compared to LB, which required increasing the Nal concentration above typical concentrations used for counterselection. These selective plates were incubated for 7-8 days at 30°C. Colonies that emerged were struck on fresh selective 0.5x marine broth agar plates and transformants were checked using the same approach described above for electroporated transformants.

### Two-step chromosomal allele replacement

A standard two-step allele replacement approach with sucrose/*sacB* counterselection (Hmelo *et al*., 2015, Ried & Collmer, 1987) was optimized for *E. litoralis* DSM 8509. pNPTS138-derived allele replacement plasmids were introduced by conjugation as described above. Chloramphenicol resistant clones were inoculated into 2 ml 0.5x marine broth and allowed to grow without selection for 8-24 hours. These cells were spread on 0.5x marine broth agar supplemented with 7.5% sucrose to select for clones in which a second recombination event excised the plasmid. After approximately 1 week of growth at 30°C, sucrose resistant colonies were replica patched on agar plates with or without chloramphenicol. From the chloramphenicol sensitive clones (i.e. clones in which the plasmid had excised) the locus of interest was PCR amplified and sequenced to distinguish clones bearing the parental allele from those bearing the desired new allele.

### PacBio whole-genome sequencing

*E. litoralis* DSM 8509 genomic DNA was extracted using guanidium thiocyanate as previously described (Pitcher *et al*., 1989). Standard Pacific Biosciences (PacBio) large insert library preparation was performed. Briefly, DNA was fragmented to approximately 20kb using Covaris G tubes. Fragmented DNA was enzymatically repaired and ligated to a PacBio adapter to form the SMRTbell template. Templates larger than 10kb were BluePippin (Sage Science) size selected, annealed to sequencing primer, bound to polymerase (P6), bound to PacBio Mag-Beads and SMRTcell sequenced using C4 chemistry. The genome was sequenced using two SMRT cells and was assembled de novo using HGAP3 and polished with quiver. This process yielded a single, closed contig. The genome was automatically annotated using the DOE-JGI pipeline (Huntemann *et al*., 2015). HWE/HisKA_2-family histidine kinases and the core GSR regulators *phyR*, *ecfG*, and *nepR* were identified manually and annotated before sequence submission to GenBank. Raw PacBio reads can be accessed through the NCBI sequence read archive at accession SRS1630618. The complete, annotated sequence of *E. litoralis* DSM 8509 is available at GenBank accession CP017057.

### RNA extraction

For RNA-seq and qRT-PCR, RNA was extracted using a Trizol reagent based protocol. First, cells were struck from freezer stocks onto fresh 0.5x marine broth agar plates. After 3 days of growth, a scoop of mixed colonies of cells were inoculated into 2 ml 0.5x marine broth in 13 × 100 mm culture tubes and grown for 24 hours 30°C shaking on a 45 degree incline at 200 RPM. Cultures were diluted to 0.005 OD_660_ in 2 ml fresh 0.5x marine broth and allowed to continue growth at 30°C shaking at 200 RPM. After 20-22 hours, these cultures were diluted to 0.001 OD_660_ in 6 tubes (for RNA-seq) or 2 tubes (for qRT-PCR) with 2 ml each of 0.5x marine broth. Half of the tubes were placed in the top row of a shaker rack under a bay of 10 T8 fluorescent light bulbs (light conditions). The other half were wrapped in foil (dark conditions) and grown in the bottom row of the same shaker rack. After 22-24 hours of growth in constant light or constant dark conditions, culture densities reached 0.15-0.25 OD_660_. Cells were harvested by rapid centrifugation in small “genotype-condition” batches to minimize handling time between culture growth and cell lysis. For RNA-seq, the 3 × 2 ml from each genotype-condition set were distributed over 4 × 1.5 ml microfuge tubes and centrifuged for 60-90 seconds at 14,000-17,000 × g. After quickly aspirating the supernatant liquid, the four pellets were resuspended in a total of 1 ml Trizol reagent (Invitrogen). For qRT-PCR, 1.5 ml of culture was harvested as above and resuspended in 1 ml Trizol. Cells lysed in Trizol were immediately stored at −80C until extraction. For RNA-seq, five samples of each genotype-condition were collected, each grown on a different day.

To extract RNA from the Trizol suspended cells, samples were thawed and incubated at 65°C for 10 minutes. After addition of 200 µl of chloroform, samples were vortexed, incubated on the benchtop for 5 minutes, then centrifuged for 12-15 minutes at 14,000-17,000 × g. The aqueous phase (∼500 µl) was transferred to a new 1.5 ml tube and 450 ul isopropanol was added to precipitate the nucleic acid. Samples were frozen overnight at −80°C then centrifuged at 17,000 × g for 30 minutes at 4°C. The pellet was washed twice with 750 µl cold 70% ethanol, suspended in RNase free water (200 µl for RNA-seq samples and 100 µl for qRT-PCR samples), and stored at −80°C.

### RNA-seq

RNA was subjected to DNase digestion using TURBO DNase I (Thermo Fisher). RNA samples were bound to an RNAeasy column (Qiagen) and digested on the column by application of 70 µL of DNase coctail (7 µL DNase, 7 µL 10x Buffer, 56 µL diH_2_O). Digests were incubated at room temperature for 45 minutes. RNA was cleaned and eluted from the column using buffers provided with the RNAeasy columns. DNA removal was confirmed by PCR using the RNA samples as a template and primers that amplify ∼ 100 bp products (set 1: F-GACGGAGAAAAAGGCATCGC, R-GATTCGCCGTGTTCATCTGC; set 2: F-CCCACGAACCGATTTCATGG, R-CCTTCGGGGAGTTTCAAGCA). Absence of amplification confirmed removal of contaminating genomic DNA. Amplification from pre-DNAse samples served as a positive control. rRNA was depleted from the sample using the Gram-negative bacteria Ribo-Zero rRNA Removal Kit (Illumina-Epicentre). RNA-seq libraries were prepared with an Illumina TruSeq stranded RNA kit according to manufacturer’s instructions. The libraries were sequenced on an Illumina HiSeq 4000 by the University of Chicago Functional Genomics Core Facility. Data were analyzed using the RNA-seq workflow in CLC genomics workbench v11.0. RNA-sequencing reads have been deposited in the NCBI GEO database under accession GSE126532.

### qRT-PCR

The GSR-dependent transcript, *dps* (Ga0102493_111653), and an endogenous control transcript (Ga0102493_112759) were evaluated using TaqMan probes and SuperScript III Platinum One-Step qRT-PCR Kit (Invitrogen) with a QuantStudio 5 real time PCR system (Thermo Fisher). The primer-probe sets (*dps* F – TCTCATCGCCGAACTCAAC; dps R – CGTGCCAGTGGAAATTCTTG; *dps* probe –/5HEX/CTTCGCGCT/ZEN/GTTCACCAAGACC/3IABkFQ/) and (ctrl F – AGATCGAAATGCTGTTGAAACG; ctrl R – GACCATCCAGAACGACATCC; ctrl probe - /56-FAM/CCGCAACAC/ZEN/CTATATCTACCCGCC/3IABkFQ/) were custom prepared by Integrated DNA Technologies. In control one-step RT-PCR reactions with a dilution series of template, both primer-probe sets exhibit 91-94% efficiency. Primers and probes were mixed to make a 10 µM F, 10 µM R and 5 µM probe stock solution. 0.4 µl primer-probe stock solution was used in each 20 µl reaction. 50 nM ROX served as the reference dye in each reaction. Each RNA sample was diluted to 2.5 ng µl^−1^; 4 µl were used in each 20 µl reaction. Each sample was assayed in triplicate for each probe set. The average Ct from these technical replicates was considered the Ct for the sample. “No RT” control reactions were conducted on each sample with each probe to ensure that the signal from contaminating genomic DNA was less than 5 % of the signal (i.e. that the Ct from the “No-RT” reaction was at least 4 cycles later than the Ct from the “+RT” reaction. Two control samples (WT – Dark, and WT – Light samples) were diluted to 2.5 ng µl^−1^ and frozen in 50 µl aliquots. One aliquot was thawed and assayed on each plate to ensure consistency between plates. Reaction parameters were: 50°C 4 min, 95°C 5 min, followed by 40 cycles of 95°C 15 sec, 60°C 30 sec where fluorescence was measured each cycle. For each sample, ΔCt (Ct(*dps*)-Ct(control)) was calculated. Then the average WT-dark ΔCt was subtracted from each ΔCt to generate a ΔΔCt, which is the same as log_2_(fold change) compared to WT-dark.

### Protein expression and spectroscopy

*E. litoralis lovK* (gene locus Ga0102493_111685) and a variant of *lovK* in which the flavin adduct-forming cysteine was mutated to an alanine (C73A) were cloned into the pET28a expression plasmid; see Table S6 for primers. Protein was expressed from these plasmids in *E. coli* BL21(DE3). Briefly, 2 liters of LB (containing 50 µg ml^−1^ kanamycin) was inoculated with a 100 ml of an overnight culture. The culture was grown at 37°C / 220 rpm to OD_600_ ≈0.8 and induced with 0.5 mM IPTG for four hours before cells were pelleted by centrifugation. For purification, the cell pellet was resuspended in 50 ml of resuspension buffer (10 mM Tris (pH 7.6), 150 mM NaCl, 10 mM imidizole) and lysed by two passages through a LV-1 microfluidizer. The lysate was then clarified by centrifugation (15 minutes at 35,000 × g) and applied to 2.5 ml of Ni-NTA resin on a gravity column. The resin had been pre-equilibrated with lysis buffer. The resin was washed with 5 column volumes (CV) of resuspension buffer, followed by a step gradient of buffer with increasing concentrations of imidazole. Specifically, the column was washed with 12 ml resuspension buffer containing 75 mM imidazole, 4 ml buffer with 200 mM imidazole, and finally 5 ml buffer with 500 mM imidazole for elution. The purity of the different fractions was assessed on a 12% SDS-PAGE gel. Peak fractions were desalted with a Zeba spin column (ThermoFisher) with a 7000 kDa MWCO. The visible absorption spectrum of purified LovK and LovK(C73A) was measured in a Tecan Spark 20M. The “lit” state of LovK was generated by illuminating purified protein with a panel of 96 LED bulbs (3 mm bulbs, 430 nm peak wavelength) held 10-20 cm from the sample for 30 seconds (until the visible yellow color was bleached).

## Supporting information

Table S3: RNA-Seq transcriptome analysis for all genes and all samples

Table S4: GSR regulon

Table S5: Light-dark regulon

## ACKNOWLEDGEMENTS

We dedicate this study to the memory of Winslow Briggs, whose contributions to the field of photobiology cannot be overstated. We thank members of the Crosson lab and Maureen Coleman for helpful discussions throughout this work. We thank Angie Schmoldt at the Great Lakes Genomics Center at UW-Milwaukee for support with PacBio sequencing. The authors have no conflicts of interest to declare.

## AUTHOR CONTRIBUTIONS

Conception and design of the study, AF and SC; acquisition, analysis, or interpretation of the data AF, LV, XAN and SC; writing of the manuscript, AF and SC.

