## Supplemental Tables 1,2,6 and Supplemental Figures for "Regulation of the *Erythrobacter litoralis* DSM 8509 general stress response by visible light"

---

**Table S1.** PacBio HGAP3 genome assembly statistics for *Erythrobacter litoralis* DSM 8509[1].

| | |
|---|---|
| Number of SMRT cells used | 2 |
| Number of bases sequenced | 523,219,652 |
| N50 read length (bp) | 11,755 |
| Mean read length (bp) | 7,631 |
| Mean read score | 0.86 |
| Assembled reads | 64,175 |
| Number of closed assembled replicons | 1 |
| Length (bp) of assembled replicon | 3,251,283 |
| Average Depth of Coverage | 143 |

[1] All data (raw PacBio reads, sample metadata, and assembled and annotated genome) are available through NCBI at the following accessions:

NCBI GenBank locus number CP017057
NCBI Bioproject accession number PRJNA339618
NCBI Biosample accession number SAMN05601187
NCBI sequence read archive (SRA) accession number SRS1630618

**Table S2.** Antibiotic sensitivity of DSM 8509 on 0.5 X Marine broth agar.

| **Antibiotic** | **Minimal concentration (in ug/ml) to reduce CFU by at least 4 orders of magnitude.** |
|---|---|
| Ampicillin | 50 |
| Apramycin | 100 |
| Carbenicillin | 25 |
| Chloramphenicol | 0.5-1 |
| Gentamycin | 10 |
| Kanamycin | 25 (high spontaneous resistance ~ 1 per $10^5$-$10^6$ cells) |
| Rifampicin | 5 |
| Spectinomycin | 10 (spontaneous resistance noted) |
| Streptomycin | Resistant (maximum tested 100) |
| Tetracyclin | 100 (Resistant to 50) |

**Table S3.** RNA-Seq transcriptome analysis for all genes and all samples (Excel spreadsheet)

**Table S4.** GSR regulon (Excel spreadsheet)

**Table S5.** Light-dark regulon (Excel spreadsheet)

### Table S6: Plasmids, primers and strains used

| ***E. coli* strains with plasmids** | | | |
|---|---|---|---|
| **Strain #** | **Genotype** | **Comments / primers to amplify inserted sequence** | **Source** |
| FC929 | TOP10 | Cloning strain | Invitrogen |
| FC3 | MT607 / pRK600 | Helper strain for tri-parental matings | Finan *et al.*, 1986 |
| FC803 | Rosetta(DE3) pLysS | Strain for heterologous protein expression | Novagen |
| **Expression plasmids for complementation**<br>**gene including ~300 bp upstream of the coding sequence were cloned into NdeI and XbaI sites** | | | |
| MTLS 4419 | TOP10 / pBVMCS-4 | Replicating plasmid, BBR origin, GentR | Thanbichler *et al.*, 2007 |
| FC3306 | TOP10 / pBVMCS4-$P_{nepR}$-*nepR-ecfG* | F: ccaacatatgGAGGAAGATCGCGACGGGAAT<br>R: tggttctagaCTTTTTCATGTGCGGCGATGTTTTC | This work |
| FC3307 | TOP10 / pBVMCS4-$P_{phyR}$-*phyR* | F: attccatatgCTCGGGATCGTATCACCAGC<br>R: attctctagaCTTCAGCCGTCCTCGAGC | This work |
| FC3308 | TOP10 / pBVMCS4-$P_{gsrP}$-*gsrP* | F: tggtcatatgCCGCCATTTGTGGTTCCCATG<br>R: tggttctagaGGCGCTTTTCTTCGGATCGAG | This work |
| FC3309 | TOP10 / pBVMCS4-$P_{gsrK}$ *gsrK* | F: tggtcatatgGGTCGTGTCGCGGATCATCTC<br>R: tggttctagaTAACACGCGCACGAGCCTAGAG | This work |
| FC3310 | TOP10 / pBVMCS4-$P_{lovK}$-*lovK-lovR* | F: tattcatatCGAATGCGTCGCAGGAAAAA<br>R: tatttctagaATCCTGCGCAGTTCGAGATT | This work |
| **Allele replacement plasmids**<br>**Inserts generated by overlap extension PCR, digested, and ligated into appropriately digested plasmid** | | | |
| FC 2443 | TOP10 / pNPTS138-CAT | Allele replacement plasmid, ChlorR, SacB | Herrou *et al.*, 2018 |
| FC3311 | TOP10 / pNPTS-CAT-$\Delta$*nepR-ecfG* | UP F: ttgcctgcagATAGGACGAAAAACTGGTTC<br>UP R+: ATTGAGCG CTGCTTCTCTTGATGTTTCCTGTGAC<br>DN F+: GAAGCAG CGCTCAATTCCATGCTCGCCGATGG<br>DN R: caataagcttCAGACCTGGAATCACGCCTTCTACTGGAA | This work |
| FC3312 | TOP10 / pNPTS138-CAT-$\Delta$*phyR* | UP F: tattctgcagCCAGCTCGCCTTCATGATGA<br>UP R+: GAAGAAAAGCGC CTCGCTCAGGGACATGTAT<br>DN F+: CTGAGCGAG GCGCTTTTCTTCGGATCGAG<br>DN R: tattaagcttCATGACCTGCTCACTGGGAC | This work |
| FC3313 | TOP10 / pNPTS138-CAT-$\Delta$*gsrP* | UP F: attcggatccTTCTCGGCCATTATGCTCCG<br>UP R+: GCGCATCAC ATGTGCGCCTGAAGAGTTCA<br>DN F+: GGCGCACAT GTGATGCGCGTGCCGATC<br>DN R: atttaagcttTCATGGATCTCGAAACCGGC | This work |
| FC3314 | TOP10 / pNPTS138-CAT-$\Delta$*gsrK* | UP F: atccagatatcctgcagTCTGGTTGAGGCACAGGTTG<br>UP R+: CGATCTGTT CGAATTCGAAGCTCGCAAGG<br>DN F+: GCTTCGAATT CGAACAGATCGCACGCTAAA<br>DN R: ggctggcgccaagcttGAGGGTGTTTCTTGCATGCG | This work |
| FC3315 | TOP10 / pNPTS138-CAT-$\Delta$*lovKR* | UP F: tattctgcagGTTCAATTTCGTGGTCGGCC<br>UP R+: GCCTCGGT CCGAGAAACGATCCCCCGTCTC<br>DN F+: CGTTTCTCG GACCGAGGCGAACCGATC<br>DN R: tattaagcttAATCCGCCGACAACGAAAC | This work |
| FC3316 | TOP10 / pNPTS138-CAT-*lovK*(C73A) | UP F: tattggatcc GTTCAATTTCGTGGTCGGCC<br>UP R+: AGCGGgcGTTCCGGCCGAGGACCT<br>DN F+: CCGGAACgcCCGCTTCCTGCAG<br>DN R: cgccaagctt TCGAGGAGGAGGATCTTGCA | This work |
| FC3317 | TOP10 / pNPTS138-CAT-*lovK*(H161A) | UP F: tattggatcc GTTCAATTTCGTGGTCGGCC<br>UP R+: TCATCCGGgcCGAAAGCTCGCGC<br>DN F+: CTTTCGgcCCGGATGAAGAACA<br>DN R: cgccaagctt TCGAGGAGGAGGATCTTGCA | This work |
| **Heterologous protein expression**<br>**Plasmids were generated in TOP10 strains and transferred to Rosetta(DE3) pLysS strains for expression** | | | |
| FC155 | DH10B / pET28a | Plasmid for heterologous expression of 6-his tagged proteins from a T7 promoter, KanR. | Novagen |
| FC3318 | TOP10 / pET28-lovK | F: ttcgcatatgCGCTTCGACGACGCGCC<br>R: atttaagcttCAGACACCGGCCATCAGATC | This work |
| FC3319 | TOP10 / pET28-lovK(C73A) | Same primers as above using pNPTS138-CAT-*lovK*(C73A) plasmid as a template. | This work |
| | ***Erythrobacter litoralis DSM 8509* strains** | | **Source** |
| FC578 | Wild type *Erythrobacter litoralis* DSM 8509 (ATCC 700002) | | Yurkov *et al.*, 1994 |

| | | |
|---|---|---|
| FC3320 | DSM8509 Δ*nepR-ecfG* | This work |
| FC3321 | DSM8509 Δ*phyR* | This work |
| FC3322 | DSM8509 Δ*gsrP* | This work |
| FC3323 | DSM8509 Δ*gsrK* | This work |
| FC3324 | DSM8509 Δ*lovKR* | This work |
| FC3325 | DSM8509 Δ*gsrK* Δ*lovKR* | This work |
| FC3326 | DSM8509 Δ*gsrP* Δ*lovKR* | This work |
| FC3327 | DSM8509 Δ*gsrK* Δ*gsrP* | This work |
| FC3328 | DSM8509 Δ*gsrK* Δ*gsrP lovK(C73A)* | This work |
| FC3329 | DSM8509 Δ*gsrK* Δ*gsrP lovK(H161A)* | This work |
| FC3330 | DSM8509 Δ*gsrK* Δ*gsrP* Δ*lovKR* | This work |
| FC3331 | DSM 8509 / pBVMCS4 | This work |
| FC3332 | DSM 8509 Δ*nepR-ecfG* / pBVMCS4 | This work |
| FC3333 | DSM 8509 Δ*nepR-ecfG* / pBVMCS4-*nepR-ecfG* | This work |
| FC3334 | DSM 8509 Δ*phyR* / pBVMCS4 | This work |
| FC3335 | DSM 8509 Δ*phyR* / pBVMCS4-*phyR* | This work |
| FC3336 | DSM 8509 Δ*gsrP* / pBVMCS4 | This work |
| FC3337 | DSM 8509 Δ*gsrP* / pBVMCS4-*gsrP* | This work |
| FC3338 | DSM 8509 Δ*gsrK* / pBVMCS4 | This work |
| FC3339 | DSM 8509 Δ*gsrK* / pBVMCS4-*gsrK* | This work |
| FC3340 | DSM 8509 Δ*lovKR* / pBVMCS4 | This work |
| FC3341 | DSM 8509 Δ*lovKR* / pBVMCS4-*lovKR* | This work |


Finan, T.M., Kunkel, B., De Vos, G.F., and Signer, E.R. (1986) Second symbiotic megaplasmid in Rhizobium meliloti carrying exopolysaccharide and thiamine synthesis genes. *J Bacteriol* **167**: 66-72.

Herrou, J., Willett, J.W., Fiebig, A., Varesio, L.M., Czyz, D.M., Cheng, J.X., Ultee, E., Briegel, A., Bigelow, L., Babnigg, G., Kim, Y., and Crosson, S. (2018) Periplasmic protein EipA determines envelope stress resistance and virulence in Brucella abortus. *Mol Microbiol*.

Thanbichler, M., Iniesta, A.A., and Shapiro, L. (2007) A comprehensive set of plasmids for vanillate- and xylose-inducible gene expression in Caulobacter crescentus. *Nucleic Acids Res* **35**: e137.

Yurkov, V., Stackebrandt, E., Holmes, A., Fuerst, J.A., Hugenholtz, P., Golecki, J., Gad'on, N., Gorlenko, V.M., Kompantseva, E.I., and Drews, G. (1994) Phylogenetic positions of novel aerobic, bacteriochlorophyll a-containing bacteria and description of Roseococcus thiosulfatophilus gen. nov., sp. nov., Erythromicrobium ramosum gen. nov., sp. nov., and Erythrobacter litoralis sp. nov. *Int J Syst Bacteriol* **44**: 427-434.

### SUPPLEMENTAL FIGURES

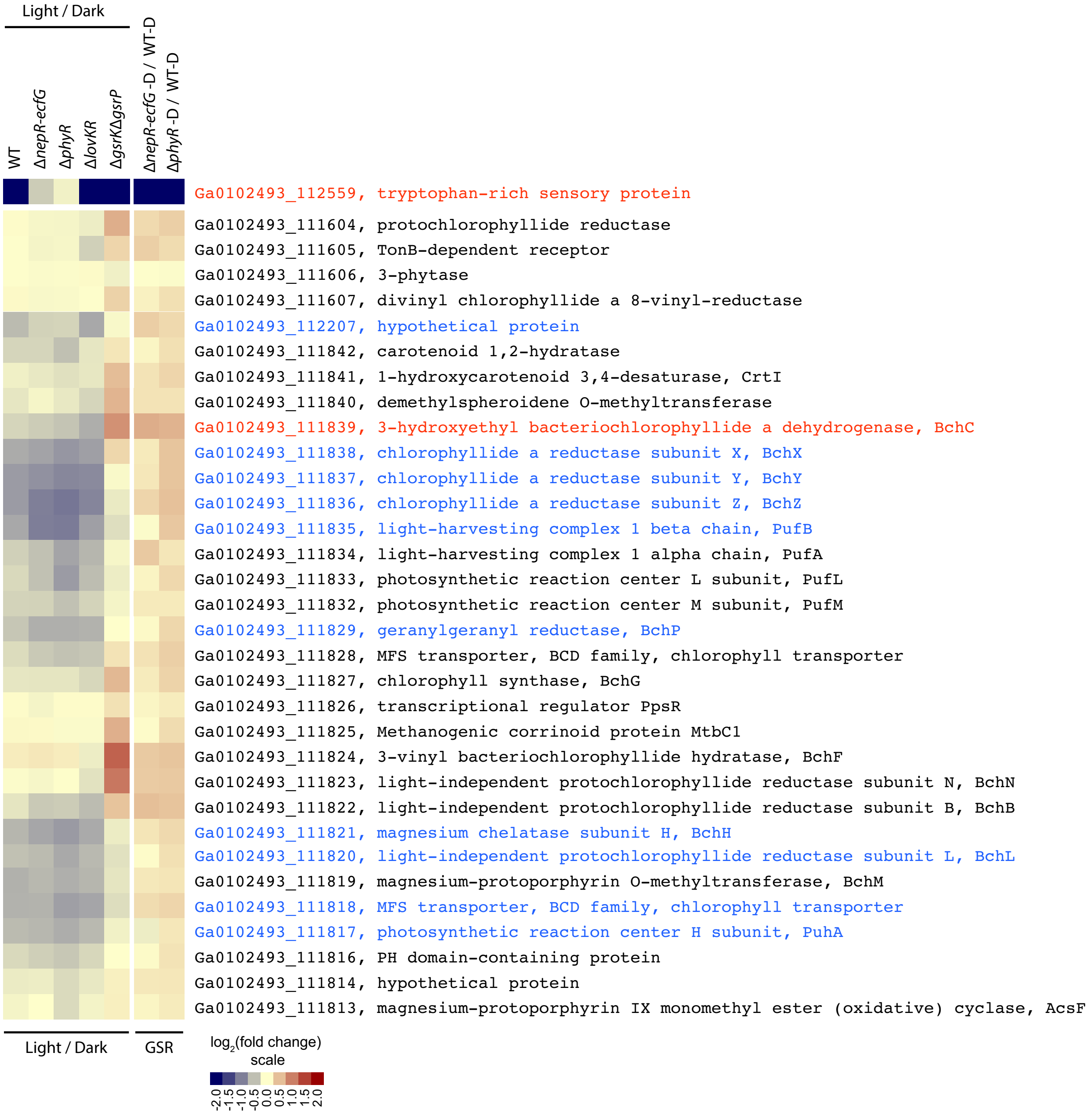


**Figure S1: Relative expression of genes involved in phototropy.**
Heatmap represents the same comparisons and same color scaling as in Figure 5. Genes were selected based on annotated functions in bacteriochlorophyll synthesis, phototrophy, or proximity to such genes. Genes are ordered by position on the chromosome. Genes in red text meet the cutoff criteria for inclusion in the GSR regulon (Table S4). Genes in blue text meet the cutoff criteria for inclusion in the light-dark regulon (Table S5).

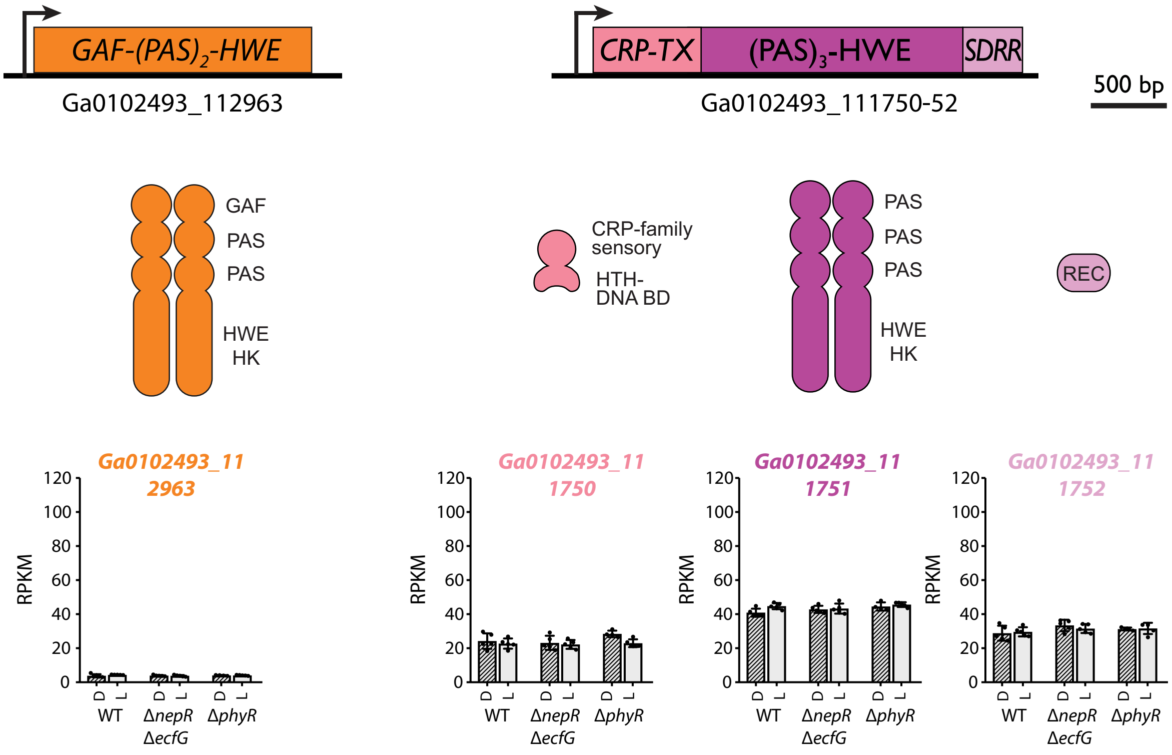


**Figure S2: Gene structure, protein domain structure and RNA-seq expression values for two additional unnamed HWE kinases encoded in the DSM 8509 genome.**

Features are drawn as in Figure 2.

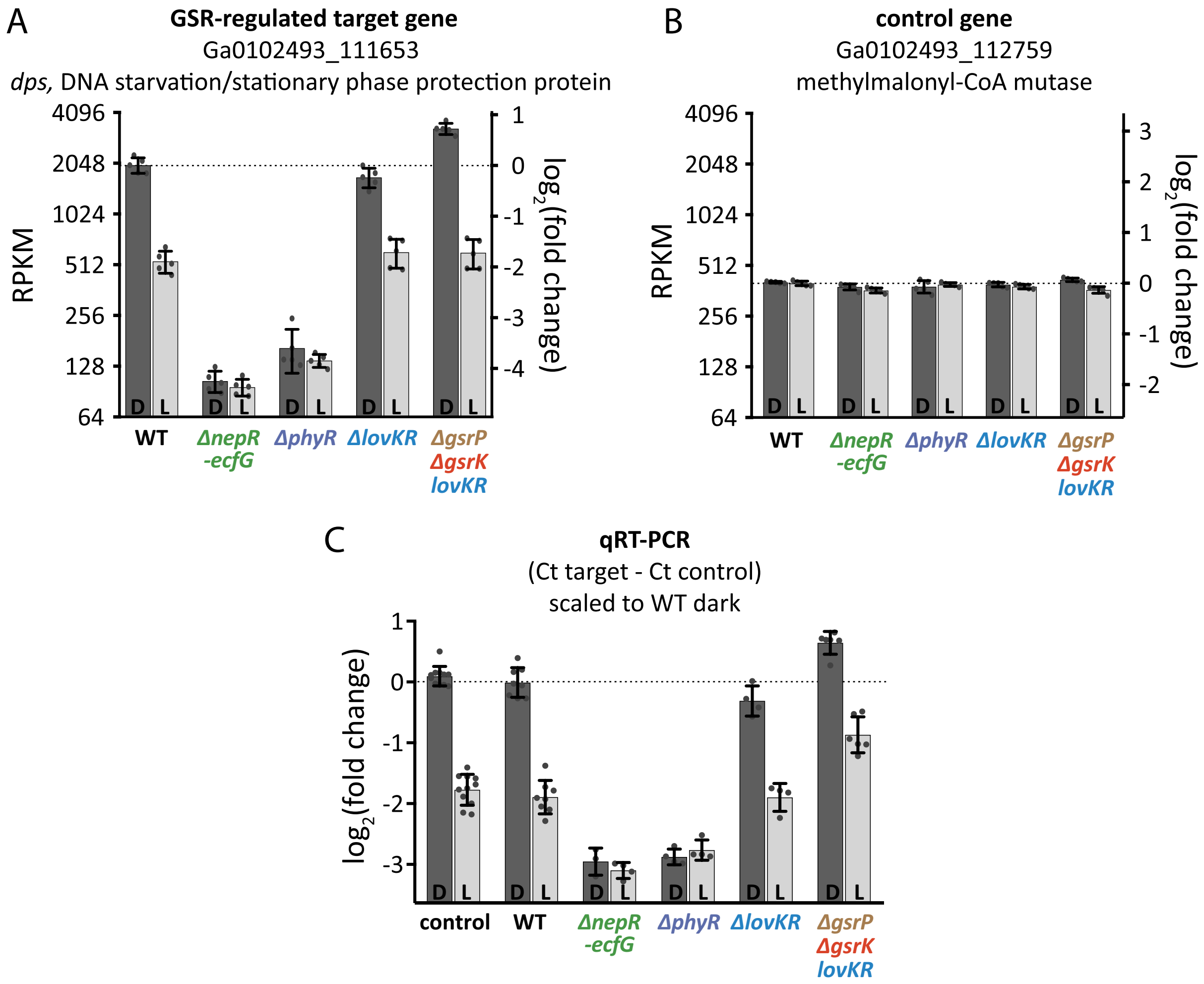


**Figure S3: Target and control genes used for qRT-PCR analysis of GSR transcription**

**A.** RPKM values for *dps* (a GSR-regulated gene), extracted from RNA-seq experiments (Table S3) plotted on a $\log_2$ scale. Bars represent mean ± s.d. of 5 independent samples (dots). $\text{Log}_2$(fold change) relative to wild type (WT) dark is scaled on the right y-axis.

**B.** RPKM values for the methylmalonyl-CoA mutase gene used as the endogenous control gene for normalization plotted as in (A).

**C.** qRT-PCR analysis of the same genotype-condition combinations assayed by RNA-seq in (A) and (B). These data are extracted from the same set of experiments presented in Figure 6B and are presented here for direct comparison to panel (A). As in Figure 6, each measurement represents the $(Ct_{dps} - Ct_{control})_{sample}$ – average $(Ct_{dps} - Ct_{control})_{WT\text{-}D}$. This results in a $\log_2$(fold change) compared to WT-dark. Strains were grown and assayed together. Each sample was assayed in triplicate. Points represent the average value for each sample. Bars represent mean ± s.d. of the samples in each condition. As an internal control, the same wild type-dark and wild type-light samples were assayed on every plate (control).

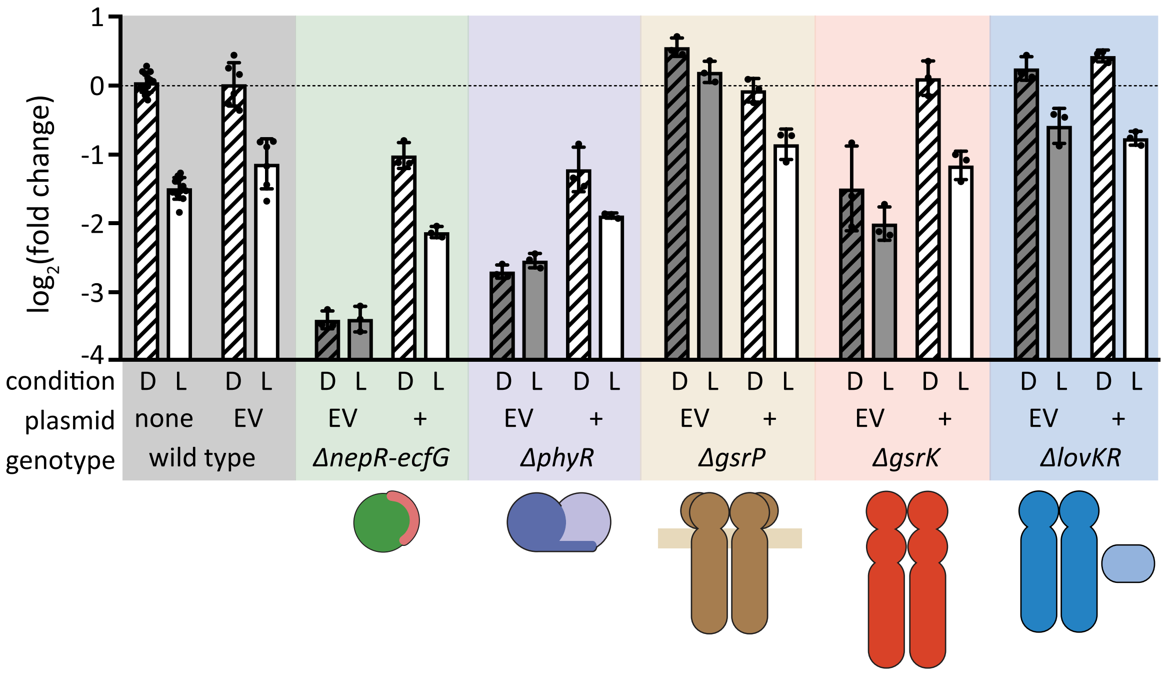


**Figure S4: Complementation of GSR transcription defects in strains with single deletions of GSR regulators.**
qRT-PCR analysis of *dps* expression as a measurement of GSR transcription. All strains, including wild type (WT), carry a replicating plasmid and were grown in the presence of gentamycin to select for plasmid maintenance. Plasmids either carried the deleted gene under the control of its endogenous promoter (+) or were empty vectors (EV) to control for plasmid and selection effects. Grey shaded bars highlight mutant strains carrying the empty vector. The chromosomal genotype of the strains in each colored block is indicated at the bottom. All strains were grown in the dark (D – striped bars) or in the light (L – open bars). Data are presented as in Figure 6 and Figure S3C. The internal control RNA samples assayed on every plate in this experiment were from wild type samples that did not carry a plasmid (far left bars).
